# Probing interplays between human XBP1u translational arrest peptide and 80S ribosome

**DOI:** 10.1101/2021.06.22.449432

**Authors:** Francesco Di Palma, Sergio Decherchi, Fátima Pardo-Avila, Sauro Succi, Michael Levitt, Gunnar von Heijne, Andrea Cavalli

## Abstract

The ribosome stalling mechanism is a crucial biological process; yet its atomistic underpinning is still elusive. In this framework, the XBP1u translational arrest peptide (AP) plays a central role in regulating the Unfolded Protein Response (UPR) in eukaryotic cells. Here, we report multi-microseconds all atom molecular dynamics simulations designed to probe the interactions between the XBP1u AP and the mammalian ribosome exit tunnel, both for the wild type AP and for four mutant variants of different arrest potency. Enhanced sampling simulations allow investigating the AP release process of the different variants shedding light on this complex mechanism. The present outcomes are in qualitative/quantitative agreement with available experimental data. In conclusion, we provide an unprecedented atomistic picture of this biological process and clear-cut insights into the key AP-ribosome interactions.

## Introduction

Far from being an inert conduit for the nascent polypeptide chain (NC), the ribosome exit tunnel provides an environment with many opportunities for NC residues to form interactions of different degrees of stability with rRNA or ribosomal proteins in the tunnel wall. A particularly striking example of biologically relevant NC-exit tunnel interactions is provided by the so-called translational arrest peptides (APs), relatively short stretches of a polypeptide that have evolved to stably interact with the exit tunnel in such a way that the geometry of the ribosome active site – the polypeptide transferase center (PTC) – is sufficiently distorted to block further elongation of the NC^1^. In many cases, the elongation arrest can be relieved by an external force “pulling” on the NC. As a result, APs act as natural force sensors in various regulatory systems in prokaryotic and eukaryotic cells. AP-based force sensors can also be used as an experimental tool to study co-translational processes such as protein folding, protein translocation, or insertion of proteins into membranes^2–4^.

Eukaryotic cells have evolved a sophisticated regulatory system to alleviate endoplasmic reticulum (ER) stress caused by the accumulation of misfolded proteins in the lumen of the ER: the unfolded protein response (UPR)^5^. The IRE1 sensor controls one branch of the UPR in the ER membrane, and, when activated, splices the XBP1u mRNA to generate a frame-shifted version that codes for the nuclear transcription factor XBP1s. XBP1s, in turn, activates the transcription of genes encoding protective ER-stress proteins^6,7^. The XBP1u mRNA is recruited to the ER membrane in the vicinity of IRE1 by virtue of a moderately hydrophobic segment in the XBP1u protein that binds to the Sec61 translocon. An AP immediately downstream of the hydrophobic segment stalls translation of XBP1u, giving rise to arrested, Sec61-bound mRNA-ribosome-NC complexes primed for splicing by activated IRE1 and production of the XBP1s transcription factor^8,9^. The XBP1u AP thus has a central role in the UPR. Interestingly, an extensive mutagenesis analysis of the XBP1u AP suggests that it is under selection to maintain only a moderate stalling strength. Many mutations were found that give rise to versions of the AP that require much stronger pulling forces to relieve the translational arrest^10^. Thus, it is of considerable interest to gain a detailed understanding the ribosome-AP interactions that underlie the function of the XBP1u AP.

Here, we use molecular dynamics (MD, 1 µs) and extensive replica-enhanced sampling simulations (i.e., Adiabatic Bias Molecular Dynamics (ABMD)^11^ for a total of 10 µs) to analyze XBP1u AP-mediated ribosome stalling. Starting from the structural and mutational analysis by Shanmuganathan and co-workers^10^, we study the AP-ribosome interactions for AP variants of different stalling strengths, providing new atomistic details about the XBP1u arrest mechanism in the ribosome. When subjected to external adaptive moving restraints (i.e., ABMD), we find that the XBP1u AP residues are dislodged in sequel from their respective interaction sites in the exit tunnel, starting with the most N-terminal one located near the tunnel exit portal. Additionally, we discover that specific residues in the AP, identified by earlier mutational analyses to be critical for arrest, take longer to dislodge from the exit tunnel than other residues. Finally, we follow the extraction of the AP from the exit tunnel using the ABMD protocol to better understand how a polypeptide in transit interacts with the tunnel.

## Results

The present analysis aims to unveil the dynamics of the stalled XBP1u AP in the ribosome and characterize, qualitatively and quantitatively, the stall-release mechanisms for different AP variants. Starting from the recently published cryo-EM structure^10^, we first simulated a eukaryotic 80S ribosome with the XBP1u AP stalled in the exit tunnel over 1 μs of MD simulation^12^. This allowed us to assess the complex’s thermal stability and equilibrate the system for subsequent enhanced sampling studies. Then, we ran several ABMD simulations to accelerate the dislodging and subsequent extraction of the AP from the tunnel and to investigate, at an atomistic level, how different residues in the AP influence the release kinetics. ABMD is an enhanced sampling method to accelerate rare events. In ABMD, a harmonic restraint (i.e., the bias), whose center is ruled by random thermal fluctuations occurring at room temperature, gently brings the system to the desired point in space. Thus, the ABMD method can be seen as a pawl-and-ratchet mechanical system, where the simulation temperature rules rotations.

The systems’ overall size amounts to 2.5 million atoms for the plain MD simulations and 3.0 million atoms for the ABMD simulations (see Methods). The system comprises several components: rRNAs, P-and E-site tRNAs, mRNA (6 bases, 3’-UUAAUG-5’ corresponding to Leu259 and Met260 in the XBP1u sequence), ribosomal proteins, the XBP1u AP, water molecules, and neutralizing ions. The simulated complex is shown schematically in Fig. 1, highlighting the main molecular entities in play.

**Figure 1.**
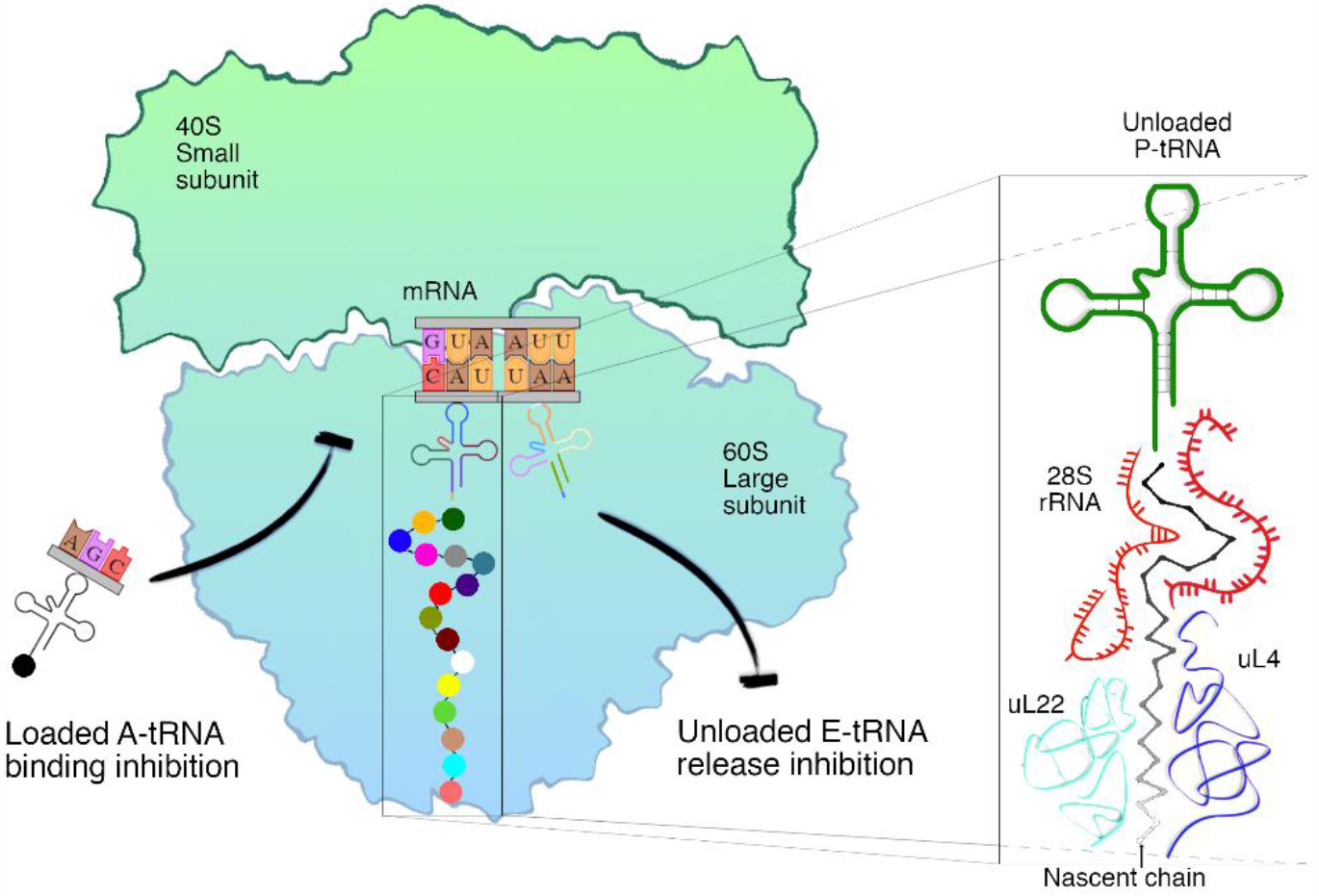
The paused eukaryotic ribosome in post-state (PDB id: 6R5Q), with incoming A-site tRNA, with the AP (in balls in the exit tunnel), the mRNA, the P-tRNA (green in the right inset) and E-tRNA. In the inset the P-tRNA (green), 28S rRNA (red) and uL22 (cyan) and uL4 (blue) proteins are shown in detail together with the nascent chain in a black/grey gradient.

To relax the structure before MD simulations, the ribosome was initially energy-minimized, then solvated, again minimized, and finally slowly thermalized and pressurized during the equilibration step (see Methods). In the following, we report on the MD and the ABMD simulations.

### The stalled ribosome

This first step in our analysis aimed to understand the system’s overall behavior through unbiased (plain) MD simulations, i.e., without any acceleration of rare events to overcome free energy barriers. In the following, whenever referring to residues in the AP, we prepend one of the three AP portions as defined in Fig. 2a to the residue name: the C-terminal (CT-) portion of the AP (residues 254-260), the intermediate (I-) portion (residues 247-253), and the N-terminal (NT-) portion (237-246). For the ribosome, we prepend the subunit name to the name of the residue or nucleobase.

**Figure 2.**
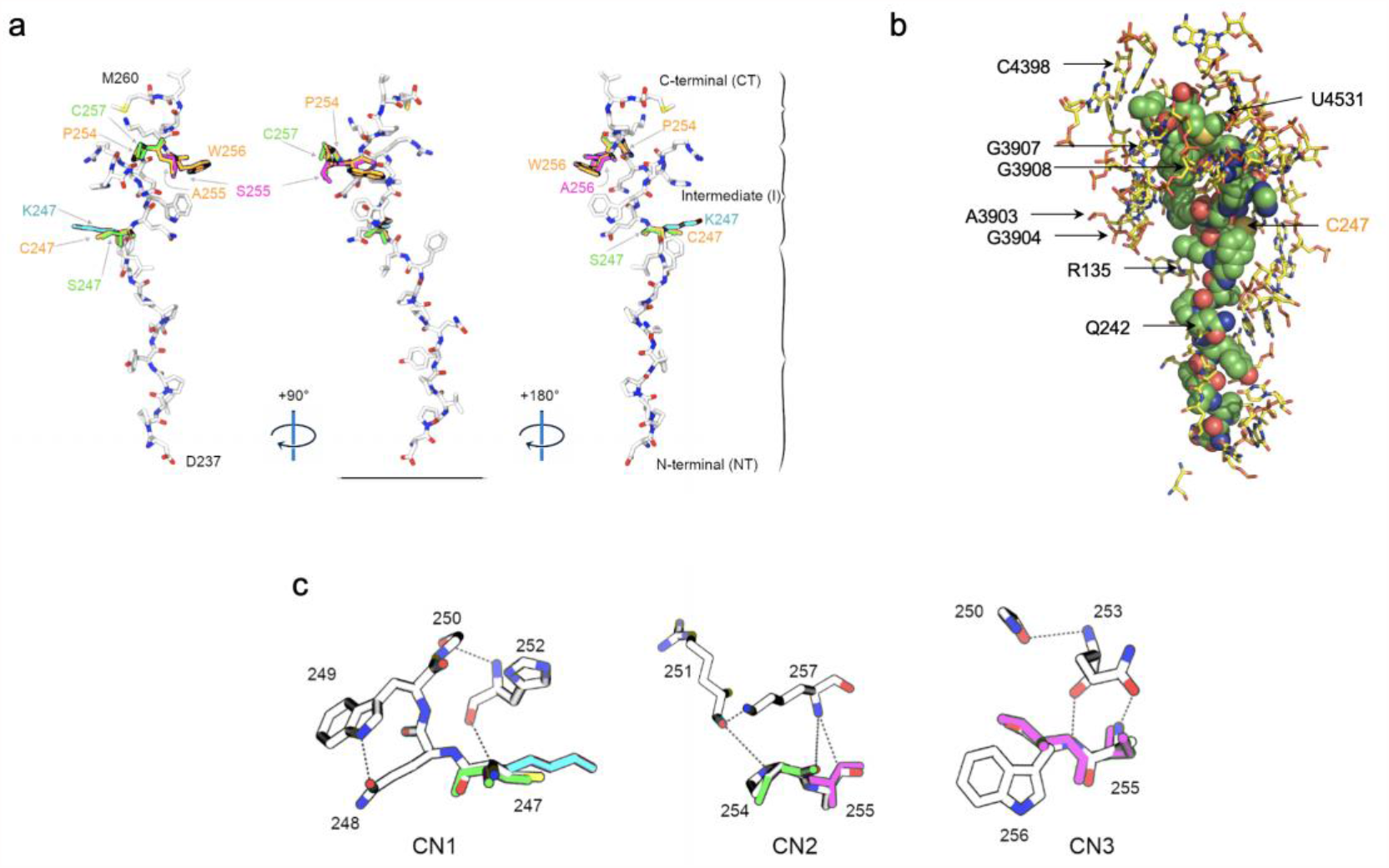
a) The human XBP1u[S255A] AP from three different view-points (image trio with stereo angles of 90° and 180°) as it is found in the CryoEM structure inside the ribosomal exit tunnel. For clarity the AP is split in three portions (braces on the right): a C-terminal portion (CT, residues 254-260), an intermediate portion (I, residues 253-247) and an N-terminal portion (NT, residues 237-246). The 24 amino acids in the AP are shown in licorice representation colored by atom name (C in white, N in blue, O in red and S in yellow). The mutated residues (247, 254, 255 and 256) in the different variants are overlayed, and labelled in orange (S255A), magenta (W256A), green (C247S/P254A/S255A) and cyan (C247K/S255A). The N-and C-terminal residues (D237, M260) are indicated. b) Ribosome residues/nucleobases (in yellow sticks) within 3.5 Å of the AP (in green spheres) after the 1 μs all-atom MD simulation. Residues discussed in the text are indicated. C247 in the AP is indicated for reference. c) Main intramolecular contacts (CN1, CN2, CN3) maintained during the simulation; same color coding as in panel a.

During the entire 1 μs MD simulation, the RMSD of the tRNAs, the mRNA, and the AP is low, confirming that the system is in a stable configuration (Supplementary Fig. 1). In particular, the AP showed a lower RMSD than the whole ribosome. Additional supporting evidence of the stability of the arrested state is the persistency of the interaction between ribosomal amino acids/RNA bases lining the exit tunnel and AP residues. Indeed, all the prominent AP-ribosome and AP-AP interactions identified in the cryo-EM structure^10^ are stable during the MD simulation (Supplementary Tables 1 and 2).

**Table 1.** Observables from the 20 100ns-long ABMD replicas for the WT and four XBP1u AP variants. The computed observables are compared with fraction full-length protein as measured in References 9 and 10.

| AP variant | Met260 detach time from PTC ($\langle t \rangle_{M260}$) | AP extraction Time ($\langle T \rangle$) | #Detached replicas | #Extracted replicas | Covered distance (D) in CV space | Experimental Values ($f_i$) |
| --- | --- | --- | --- | --- | --- | --- |
| WT | 13.2 ns ± 0.4 | 47.2 ns ± 1.3 | 20/20 | 16/16 | 12.5 nm ± 0.00 | 1.0 |
| S255A | 14.0 ns ± 0.9 | 47.9 ns ± 1.6 | 20/20 | 20/20 | 12.5 nm ± 0.00 | 0.89 |
| W256A | 13.0 ns ± 1.1 | 78.1 ns ± 3.4 | 20/20 | 15/20 | 11.72 nm ± 0.78 | 1.0 |
| C247K/S255A | No detach | No extraction | 17/20 | 0/20 | 0.47 nm ± 0.01 | 0.44 |
| C247S/P254C/S255A | 27.1 ns ± 5.6 | No extraction | 0/20 | 0/20 | 7.43 nm ± 2.65 | 0.14 |

We analyzed the molecular interactions in detail by investigating all the contacts in the minimized conformation within 3.5 Å between any atom of the AP and the ribosome (Fig. 2b; Supplementary Table 1. In particular, in the upper, C-terminal region of the AP (near 28S rRNA and P-tRNA), we found CT-Leu259 stably interacting with 28S-C4398; this cytosine is engaged in an interaction with the isobutyl side chain of the leucine, thus stabilizing a conformation that would clash with an incoming acylated tRNA. As reported by Shanmuganathan et al.^10^, this is a critical contact for the silencing of the peptide transferase activity by the XBP1u AP. It is also worth mentioning the CT-Pro258 backbone oxygen in contact with the nitrogen of 28S-U4531. This nucleobase is a crucial hallmark, as it establishes a tight network of interactions with CT-Lys257, CT-Leu259, and CT-Met260. Furthermore, CT-Pro258 is surrounded by 28S-G3907 and 28S-A3908 (Supplementary Fig. 2c). Interestingly, the position and the network of interactions established by CT-Trp256 and I-Trp249 completely displaced 28S-A3903 and 28S-G3904 during the simulation. Here too, the finding is consistent with the observation by Shanmuganathan et al., who found that 28A-G3904 is partially ejected from its normal position in the cryo-EM structure^10^. As a consequence of this displacement at the beginning of the simulation (see Supplementary Table 1), 28S-U4556 becomes perfectly stacked with CT-Trp256. Furthermore, both 28S-G4551 and 28S-U4557 locked this tryptophan during the whole simulation (Supplementary Fig. 3a).

**Figure 3.**
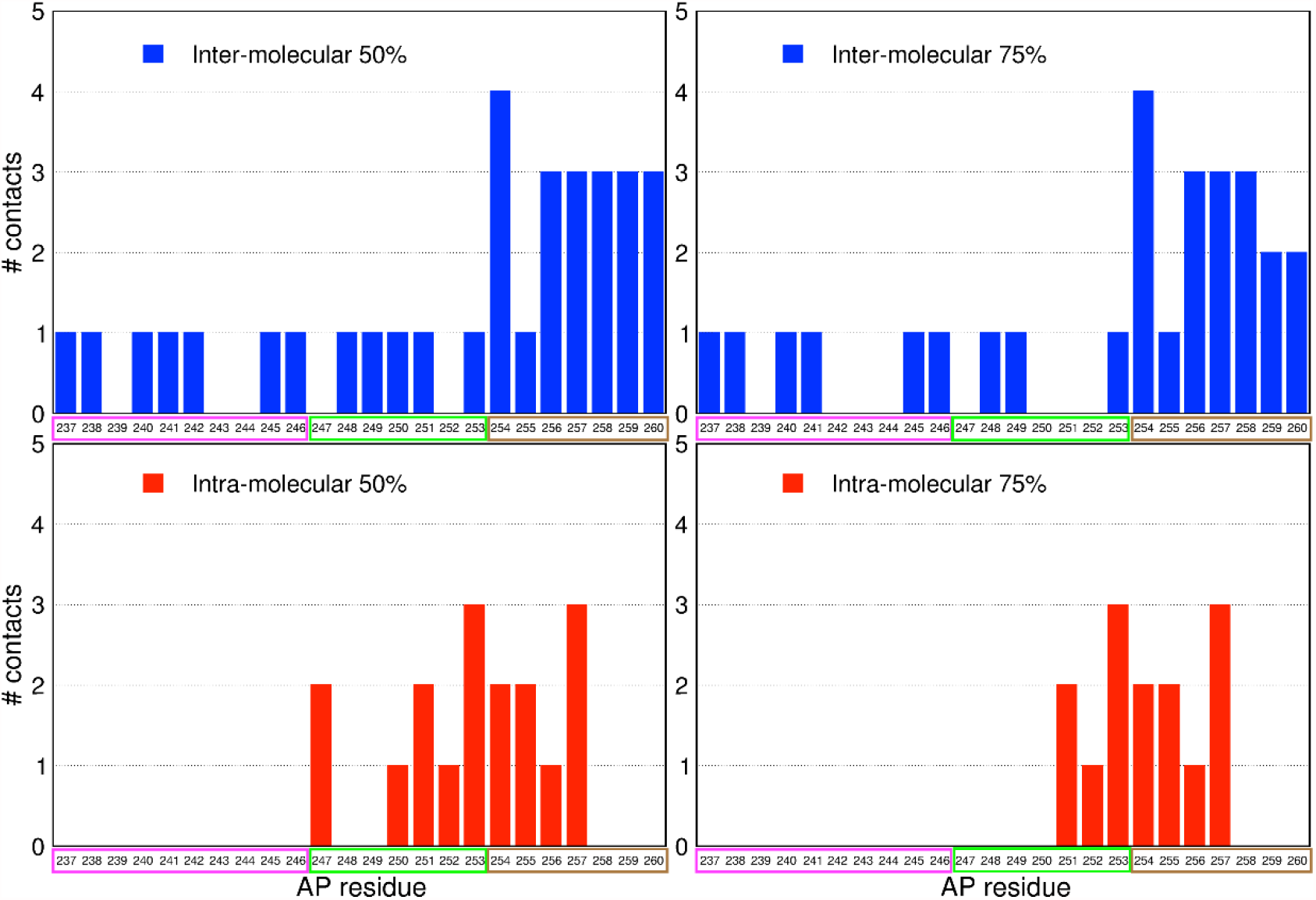
Number of intermolecular (first row) and intramolecular (second row) interactions during the 1 μs MD simulation. The interactions are counted only if present along the simulation with a persistency of at least of 50% and 75% (first column and second column respectively). The AP residues on the x-axis are framed accordingly to the NT-, I-and CT-portions, respectively in magenta, green and brown.

In the lower part of the tunnel, we found at the beginning of the simulation NT-Gln242 interacting tightly with both uL4-Gly82 and uL4-Ser87; however, after about 400 ns, these residues were replaced by uL22-Gly134 and uL22-Arg135, respectively (Supplementary Fig. 4d). Overall, the CT-region of the AP is much less dynamic than the I-and NT-regions.

**Figure 4.**
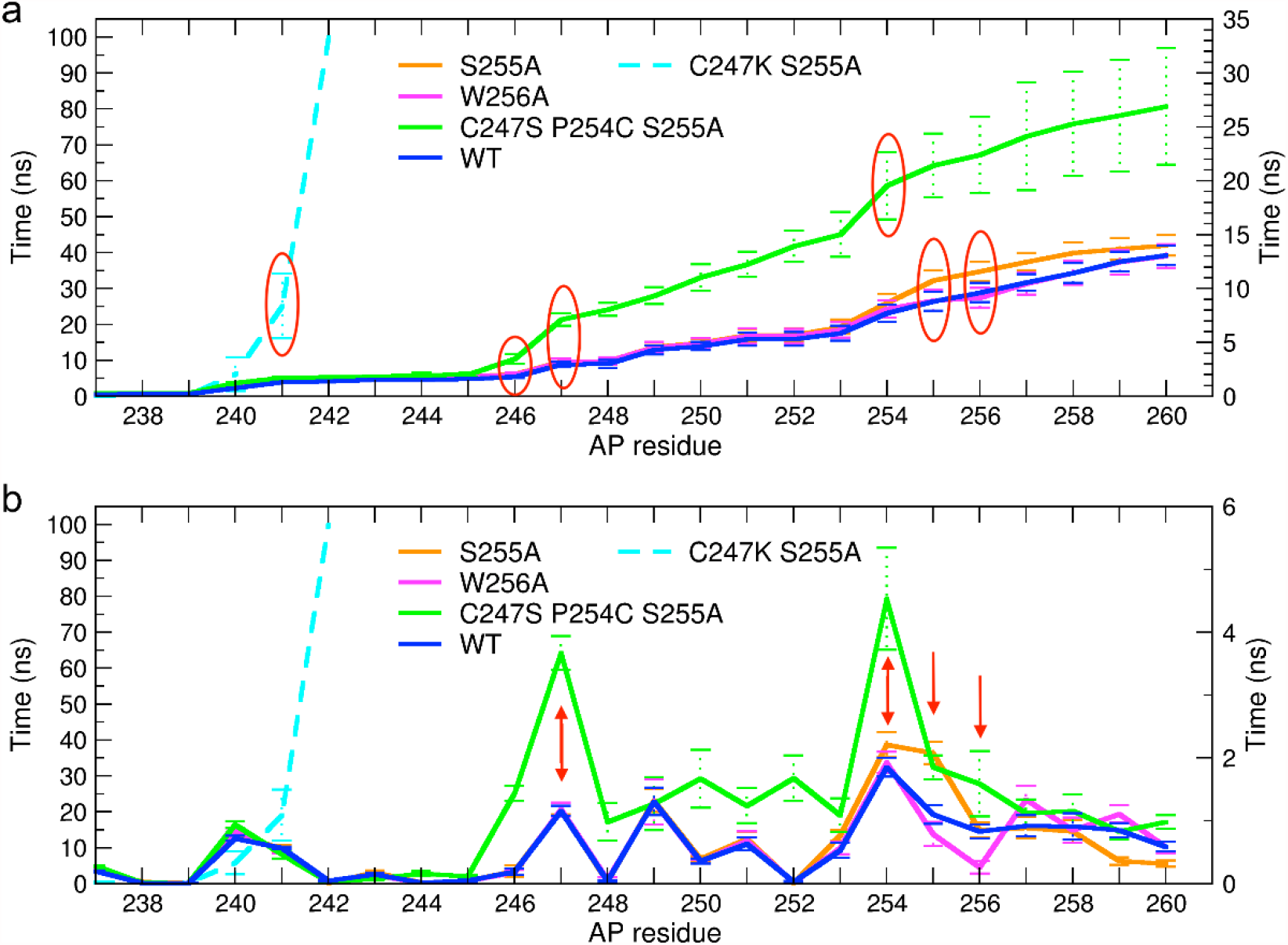
a) Average time <*t*> (and associated standard error) for the detachment of AP residues (including Leu259) from their starting position (Supplementary Table 3). b) Average time intervals <Δ*t*^*i*^ > (and associated standard error) between the detachment of residue *i-1* and residue *i* during the ABMD simulations (Supplementary Table 4). Residues C247, P254, A255, and W256 are indicated by red circles/arrows. In both panels the principal y-axis refers to the C247K/S255A variant (dashed cyan), whereas the alternative y-axis is referred to the others.

The stability of the AP intramolecular interactions was analyzed in the same way. We tracked the trend along the trajectory of all the interactions reported in Shanmuganathan et al.^10^ and a few additional interactions established during the simulation. For convenience, we grouped these intramolecular contacts into three interconnected networks (Fig. 2c): I-Cys247, I-Gln248, and I-Trp249 (CN1); I-Arg251, CT-Pro254, CT-Ala255, and CT-Lys257 (CN2); and I-Gly250, I-Gln253, and CT-Trp256 (CN3). According to our simulations, the most stable network was CN3 (Supplementary Fig. 5d); however, CN1 and CN2 were still contributing to the rigidity of the AP structure in the second half of the simulation, helping to maintain the ribosome in the stalled state. The interactions between I-His252 and I-Cys247/I-Gly250 stabilized after 500 ns (Supplementary Fig. 5a) and the CT-Ala255/CT-Lys257 pairing was found to fluctuate between two states. CT-Lys257 also made a relatively stable H-bond with I-Arg251, prone to fluctuate during the simulation (Supplementary Fig. 5b).

**Figure 5.**
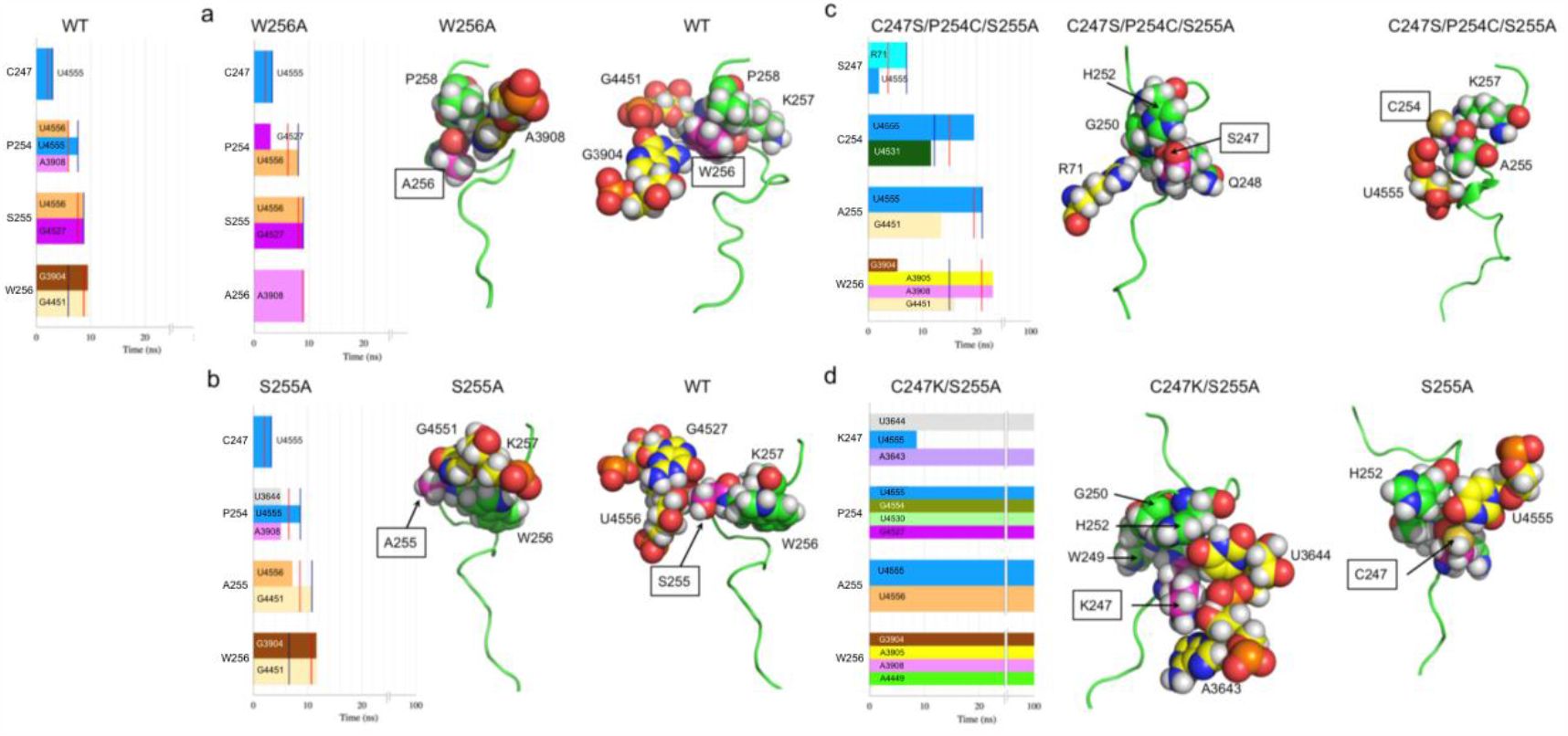
The bar graphs show conserved AP-ribosome interactions during the 100 ns ABMD simulations up to the time point *t*^*i*^ where residue *i* (*i* = 247, 254, 255, 256) detaches from its starting position in at least one of the simulations. For each residue the colored bar represents the range of time (in ns) during which the AP interacts with the specified residue/nucleobase of the ribosome. Red vertical lines indicate the time *t*^*i-1*^, where residue *i-1* detaches from its starting position, and blue vertical lines indicate the time point where residue *i* loses its last intramolecular interaction. The molecular models show the ribosome (yellow spheres) and AP (green spheres and ribbon) residues interacting with residue *i* (boxed label and magenta spheres) just before time *t*^*i*^. a) W256A, b) S255A, c) C247S/P254C/S255A, d) C247K/S255A. The WT bar graph is shown top left.

In summary, the system was very stable for 1 μs. However, some intriguing differences between the AP regions could be observed. The C-terminal region displayed a number of persistent AP-ribosome intermolecular contacts, whereas the upper-central part of the tunnel showed chiefly intramolecular AP interactions. Fig. 3 (upper panels) shows that the number of persistent intermolecular contacts in the C-terminal region is significantly higher than those in the central and N-terminal regions. This information was retrieved from the entire MD trajectory, and only the contacts present for at least 50% and 75% of the simulation time were reported. Two thresholds were used to confirm the stability of the result, an approach similar in spirit to persistence homology theory^13,14^. The same quantitative analysis was performed for intramolecular AP contacts; the results show that the upper-intermediate region is where the highest number of interactions are consistently present (Fig. 3, lower panels). Thus, many intermolecular contacts are established between the AP C-terminal residues CT-Pro258, CT-Leu259, and CT-Met260 and the PTC region (28S rRNA in particular) and between AP N-terminal residues and ribosomal uL22/uL4 proteins. In contrast, the compact turn region from I-Gly250 to CT-Lys257 engages in both inter-and intramolecular contacts.

### Release kinetics of different AP variants

In a second simulation campaign, we aimed to elucidate how differences in the inter-(AP-ribosome) and intra-molecular (AP-AP) interactions of the wild type (WT) and previously characterized variants of the XBP1u AP^9,10^ might explain their different stalling strengths. To reproduce the conditions of the experimental measurements, which were carried out with an external pulling force acting on the AP, we performed a new set of (time-bounded) simulations using the enhanced sampling method ABMD^11^ (a pawl-and-ratchet-like biasing technique; see Methods) and a multiple-replicas approach^15,16^. While only using thermal fluctuations, ABMD allows accelerating by several orders of magnitude the events of interest. Before the ABMD runs, the relevant mutations were introduced into the relaxed structure, which was then re-equilibrated for 10 ns. To gather sufficient statistics, we repeated the ABMD simulation 20 times for each AP variant. We analyzed the WT XBP1u AP and four variants: S255A (the one used to obtain the cryo-EM structure^10^), W256A, C247K/S255A, and C247S/P254C/S255A.

To characterize the overall release process, we used two partially dependent global observables (see Methods for details). The first one is the average time <*t>* required for the C-terminal AP residue CT-Leu259 to move > 3.5 Å away from the nucleobase P-tRNA-A76. We chose this metric because the removal of the CT-Leu259 side-chain from its starting position releases 28S-C4398 and allows the incoming A-tRNA to move into the PTC and translation to resume. The second observable is the number of replica simulations that lead to CT-Met260 detachment from P-tRNA-A76 within the 100 ns simulation time.

We compared the computational outcomes with the available experimental data from Yanagitani et al.^9^ and Shanmuganathan et al.^10^. These studies show that the W256A mutant is a weaker staller than the WT Xbp1u AP, which in turn is weaker than the S255A mutant. In contrast, the triple mutant C247S/P254C/S255A and the double mutant C247K/S255A both induce much stronger translational arrest than the S255A variant. Here, we correlated these observations with the time required by the AP to detach from the PTC. In the following, we report average times and the relative standard error, as we are interested in the uncertainty of the mean and not on the intrinsic variance of the out-of-equilibrium ABMD process.

As shown in Fig. 4a and Supplementary Table 3, CT-Leu259 took almost the same time to detach from the PTC in the WT and W256A variants (<*t*>_WT_ = 12.5 ns ± 0.9, <*t*>_W256A_ = 12.5 ns ± 1.1), while S255A took slightly longer (<*t*>_S255A_ = 13.7 ns ± 1.1). In contrast, C247S/P254C/S255A took twice as long (<*t*>_C247S/P254C/S255A_ = 26.1 ns ± 5.2). For C247K/S255A, it was not possible to estimate a detachment time as most of the AP remained in its original position during the entire 100 ns (Supplementary Table 3). CT-Leu259 detached from the PTC in all the 20 replicas for WT, S255A, and W256A variants, and 17 out of 20 replicas for the C247S/P254C/S255A variant. In contrast, CT-Leu259 did not detach from the PTC in any of the 20 replicas (Supplementary Fig. 6e).

To better understand the role of the individual AP residues in the release process, we performed a detailed analysis of the inter-and intra-molecular interactions for each variant during the simulations. For each residue *i* in the AP, we calculated the average release time <*t*^*i*^ >, and the finite-difference differential, <Δ*t*^*i*^ > = <*t*^*i*^ >-<*t*^*i-1*^ >, where *t*^*i*^ is the time during a simulation run when residue *i* first deviates > 3.5 Å from its starting position and Δ*t*^*i*^ is the time between the release of residue *i-1* and residue *i* from their starting positions (see Fig. 4 and Supplementary Tables 3, 4). The AP residues detach in sequel starting with NT-Asp237, Fig 4b, with some showing severalfold longer residence times than others. Hence the <Δ*t*^*i*^ > or <*t*^*i*^ > components, taken together, establish a kinetic signature or profile vector that can be associated with each APs.

The <Δ*t*> signature for the WT AP (Fig 4b) shows that NT-Pro240 and NT-Tyr241 have ∼5x longer residence times than the three N-terminal residues. Five to 20x increased residence times were observed for residues Cys247-Met260, except for Trp249 and His252 that had short residence times. Thus, almost all residues in the tightly packed turn region Trp249-Met260 are characterized by large <Δ*t*^*i*^ > values, correlating nicely with the high degree of sequence conservation of the turn region in Xbp1u homologs and its sensitivity to mutation^10^.

The W256A variant, which induces a weaker translational arrest than WT^9,10^, has a <Δ*t*> profile identical to the WT profile, except that the residence time for Trp256 is reduced by ∼70%. As shown in Fig. 5a, the W256A mutation removes a key interaction between NT-W249 and NT-Trp256, leaving an empty space that destabilizes the turn region. Consequently, Ala256 is released almost immediately once the preceding Ser255 has been released.

S255A, the variant used to obtain the cryo-EM structure, induces stronger translational arrest than WT^9,10^. Interestingly, as seen during the 10 ns equilibration run, solvation of the Ser255 side-chain in the WT AP prevents an important packing interaction between Gln253 and Trp256 seen in S255A (Supplementary Fig. 7). The immediate environment of A255 and S255 also differs in terms of the interacting ribosomal nucleobases (Fig. 5b). The <Δ*t*> profile of S255A differs from that of WT only for residue A255, with <Δ*t*^*255*^ >_S255A_ = 1.9 <Δ*t*^*255*^ >_WT_, leading to a slight increase in the global detach time <*t*>, in agreement with experimental data.

Experimentally, variant C247S/P254C/S255A has been found to have the strongest arrest-inducing potential of all known Xbp1u APs^10^. Indeed, the global detach time <*t*> for this variant is ∼2x longer than for WT (Fig. 4a) and its <Δ*t*^*i*^ > values are consistently higher than WT, W256A, and S255A from residue I-Leu246 to CT-Met260 (Fig. 4b). <Δ*t*^*247*^ > and <Δ*t*^*254*^ > are particularly high, reflecting the C247S and P254C mutations. As seen in Fig. 5c, compared to I-C247 in the S55A variant, I-S247 is stabilized by uL4-R71, and CT-C254 has a persistent interaction with U4555 (Fig. 5c). In both cases, these interactions lead to large increases in <Δ*t*^*i*^ >.

Finally, for the strongly arrest-inducing C247K/S255A variant, NT-Asp237, NT-Pro238, and NT-Val239 are the only residues that deviate significantly from their starting positions during the entire simulation (Fig. 4b, Supplementary Table 4). In two replicas, we observed that NT-Pro240 and NT-Tyr241 also lost their contacts with the ribosomal tunnel, whereas all other interactions remained stable in the 20 replicas. Since the S255A variant behaves differently, it is evident that the main culprit responsible for the dramatic increase in stalling strength is the C247K mutation. Indeed, as seen in Fig. 5d, two new interactions are established between I-C247K and the negatively charged phosphate backbone of the 28S rRNA by the formation of a salt-bridge with 28S-U3644 and a hydrogen bond with 28S-A3643. Additionally, there are new intramolecular interactions between I-Lys247, I-Gly250, and I-His252 (persistency > 95%), leading to extensive packing interactions around I-Lys247.

In summary, the average residence times <*t*^*i*^ > calculated for the different mutated residues agree well with the effects of the mutations on the experimentally measured stalling strength, indicating that the interactions uncovered by the ABMD simulations are functionally relevant.

### Extraction kinetics of different AP variants

The ABMD protocol also allowed us to follow the final extraction of the AP from the exit tunnel after the detachment of Met260 from the PTC (the last residue in the XBP1u AP, Asn261, is not present in our calculations because it was not yet attached to the nascent chain in the cryo-EM structure). For this reason, we did not include the covalent bond between Met260 and the nucleobase P-tRNA-A76 in the MD model; this did not affect the simulation results discussed above since Met260 stayed close to P-tRNA-A76 up until the detachment of Leu259. To characterize the extraction process, we recorded the average time <*T*> required to extract all the 24 amino acids of the AP out of the exit tunnel after Met260 had detached from the PTC (Table 1). We also recorded the number of replicas that led to fully solvated APs, and the average distance covered by the N-terminal residue in the AP (NT-Asp237) during the 100 ns simulation time.

Both the WT and the S255A variant completely left the tunnel in all the 20 replicas and in the shortest time (<*T*_*r*_ >_WT_ = 47.2 ns ± 1.3; <*T*_*r*_ >_S255A_ = 47.9 ns ±1.6). Using the same metric, despite its short detach time <*t*>, W256A took almost twice as long to extract (<*T*_*r*_ >_S255/W256A_ = 78.1 ns ± 3.4) and was able to fully exit the channel in 18 out of 20 runs. Conversely, neither C247S/P254C/S255A nor C247K/S255A were fully solvated after 100 ns of ABMD simulations.

We investigated their path along the exit channel to clarify whether the mutated AP residues play a role during the nascent chain extraction. The progression of the interactions established by AP residues 247, 254, 255, and 256 with the ribosome during the ABMD simulations of the four variants are reported in Supplementary Fig. 8 and discussed in the Supplementary Information. In general, the AP-ribosome interactions during the extraction process were different for each AP variant, even for the same residue. This likely results from the relatively fast movement of the AP through the exit tunnel during the extraction phase, not leaving enough time for the ribosome-AP interactions to equilibrate during the passage. Thus, each AP variant in a sense “sees” a different tunnel, characterized by different conformations of the amino acid residues/nucleobases depending on the AP-ribosome interactions at the time when Met260 detaches from the PTC.

## Discussion

We have used extensive MD and enhanced-sampling ABMD simulations to better understand the molecular interactions responsible for stalling the human XBP1u AP and four experimentally characterized variants in the ribosome exit tunnel. To the best of our knowledge, only one computational study of an entire eukaryotic 80S ribosome (from yeast) at the atomistic level has been published so far^17^. A few more simulation studies are available for the eubacterial ribosome^18–24^, including a recent extensive study of *E. coli* NC ejection times^25^ using both coarse-grained and all-atom steered MD simulations, and two very recent works investigating the interplay between the *E. coli* ribosome and the VemP and SecM APs^26,27^. The relative paucity of full-atom computational studies is mainly due to two reasons: the high computational power required to simulate such large systems (10^6^ atoms) for relevant timescales (i.e., µs) and the availability of reliable high-resolution structures. Both these limitations have recently been overcome. The first one thanks to the advent of GPU-based compute clusters and GPU-optimized molecular simulation engines, the latter through the development of single-particle cryo-EM^28^.

When no external force is applied, we find that the tightly packed C-terminal turn region of the XBP1u AP (residues 254-260) engages in stable intermolecular AP-ribosome interactions, while residues 250-257 are involved in intramolecular interactions within the AP. The N-terminal portion of the AP, residues 237-249, is much more mobile and adopts a more or less extended conformation during the simulation. These findings are broadly consistent with the compact conformation, strong sequence conservation, and sensitivity to mutation of residues 249-260^10^.

In the second set of simulations, we used ABMD to gently pull on the N-terminal end of the AP. The behavior of the AP was followed by measuring the average times, <*t*^*i*^ >, required for the C_a_ atom on residue *i* in the AP to be displaced >3.5 Å from its starting position, with the endpoint being the time when the C_a_ of Leu259 moves > 3.5 Å away from P-tRNA-A76. The latter time corresponds to when the sidechain of Leu259 detaches from the PTC, allowing A-tRNA to enter the PTC and translation to resume^10^.

Based on published data on how different point mutations in the XBP1u AP affect the release from the stalled state under an external force^10^, we chose to study four variants of the AP, together with the WT sequence. The global detach time from PTC, <*t*>, obtained from the simulations was in good agreement with the experimental data^9,10^. Our results show that <*t*>W256A ≈ <*t*>_WT_ ≤ <*t*>_S255A_ < <*t*>_C247S/P254C/S255A_ < <*t*>_C247K/S255A_ (Table 1), indicating that the simulations capture most or all of the essential AP-ribosome interactions responsible for stalling.

A residue-by-residue analysis of the release of individual residues in the AP from their resting positions in the exit tunnel further allowed us to identify those residues with the highest residence times, <Δ*t*^*i*^ >. For the WT AP, they are Pro240, Tyr241, Cys247, Trp249, Arg251, and Gln253-Met260. Most of these are highly conserved among XBP1u homologs and cannot be mutated without loss of stalling efficiency^10^, except for Cys247 and Pro254. Indeed, in the C247S/P254C/S255A variant, the release times for the mutated residues are 2-3 times longer than in the WT sequence. In variant C247K/S255A, the lysine residue binds so strongly to the tunnel RNA phosphate backbone, that no detachment is observed during the simulations. Likewise, the S255A mutation increases the release time for residue 255 by ∼2-fold, while W256A reduces the release time for residue 256 by about the same amount.

A direct comparison between eukaryotic and prokaryotic APs could be tricky due to their different mechanisms in inducing the ribosome stalling. In this context, while Kolář et al.^27^ performed equilibrium MD simulations only with a stalled *E. coli* ribosome, Zimmer et al.^26^ performed several replicas of steered MD^29^ with WT and mutated SecM and VemP APs in (reduced model) bacterial ribosomes. In contrast, we utilized a gentle enhanced sampling method (i.e., ABMD) with the 80S eukaryotic ribosome, highlighting subtle differences between the AP variants in atomistic detail. This is possible by taking advantage of the thermal fluctuations of the systems in the release of the different NCs.

As a technical note, we employed a time-bound of 100 ns for each ABMD simulation; this value was chosen to limit the overall computational burden and distinguish (together with the choice of the spring constant, see Methods) between the different APs. This was achieved by calibrating the simulation time such that the selected time (100 ns) and spring constant are an efficient combination that allows discerning differences between APs in a relatively limited amount of time.

We conclude that all atom MD simulations can capture essential aspects of AP-ribosome interaction unraveling atomistic details of the eukaryotic 80S ribosome arresting process, which had not been reported before, neither by experiments nor by numerical simulations. Moreover, when coupled with mutagenesis data, the combination of MD and enhanced sampling simulations can provide detailed molecular insights into how APs react to force load in the complex environment of the ribosomal exit tunnel. This study, ranking the *wt* AP and the variants in terms of detachment kinetics, and revealing the ribosome-nascent chain interactions underlying the function of the XBP1u arresting peptide, may pave the way to novel hypotheses and innovative analyses to shed further light on the complexity of mammalian ribosomes translation.

## Methods

The ribosome structure used for simulations is composed of 79 proteins and 7 RNA molecules for a total of 376837 atoms, including 300 Mg^2+^ and 5 Zn^2+^ structural ions. It is based on the 3 Å cryo-EM structure of the stalled ribosome in post-state in complex with the human XBP1u AP (PDB id: 6R5Q)^10^.

The amber14sb force field^30,^ including bsc0^31^ and χ_OL3_ ^32^ improvements for RNA, was selected to model the all-atom system. Simulations were performed with the Gromacs2020 MD engine^33^ running on the Franklin high-performance and 256-GPUs hardware platform (available at Fondazione Istituto Italiano di Tecnologia).

The first part of the protocol is based on the MD equilibration phase; upon addition in the dodecahedral simulation box of water solvent (TIP3P model^34^) and saline solution (0.15 M KCl and 0.07 M MgCl_2_, plus the ionic concentration (K^+^) required to neutralize the ribosome net charge) the system was overall composed of 2512119 atoms (including 708947 H_2_ O molecules, 5797 K^+^, 2536 Cl^-^, 408 Mg^2+^ and 6 Zn^2+^).

We run the following equilibration protocol to minimize, thermalize and pressurize the system before the 1 μs-long production run, and the AP variants systems before the ABMD simulations:

- 300 ps using a 1 fs time step and restraining 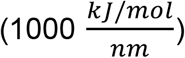 the heavy atoms of the complex during simulated annealing to bring the temperature from 0 to 300 Kelvin employing the Velocity rescale thermostat^35^ ;
- 1 ns NVT at 300 K still using a 1 fs time step and keeping the complex restrained plus 1 ns raising the time step to 2 fs;
- 1 ns NPT at 1.0 bar using the Berendsen barostat^36^ ;
- 3 steps of 1 ns each to gradually release the restrains on the complex heavy atoms (i.e., 750. 500. 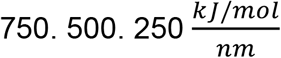) again in the NPT ensemble as in the previous step;
- 10 ns NPT without any restraint using the Parrinello-Rahman barostat^37^ to keep a 1 bar constant pressure.

The second set of simulations is based on non-equilibrium MD, specifically Adiabatic Bias MD^11^ ; in this case, we simulated five different systems, mutating some AP residues to create four variants of the *wt*: S255A, W256A, C247K/S255A, and C247S/P254C/S255A. These variants, carrying single, double or triple mutations, were chosen following the experimental evidence suggested in Yanagitani et al.^9^ and Shanmuganathan et al.^10^. The new setup of the mutated complexes in the same conditions described above was prepared by taking advantage of an experimental version of the BiKi Life Sciences software suite^38^ ; to allow the complete solvation of the NC 24 amino acids, it was necessary to break the covalent bond between the NC and the P-tRNA, and to enlarge the simulation box to include the solvent around the ribosome. This led to an increase in the total atom count to about 3 million. Hence, the systems were re-equilibrated using the same protocol described above and then simulated, adding the adiabatic bias using the PLUMED plugin version 2.7^39^ combined with Gromacs2020^33^. ABMD can evolve a system towards a target value in collective (CV) space using a harmonic potential moving along with the thermal fluctuations of the CV. This biasing potential is zero when the system moves towards the desired arrival point and damps down the fluctuations in the opposite direction. This biased MD protocol is particularly appealing and “gentle”, particularly relative to more common enhanced sampling approaches. It never explicitly pushes the system towards the desired direction but prevents the system from going back in CV space as in the pawl and ratchet mechanical system. This is particularly advantageous compared to other approaches, like steered MD^29^ (or other methods), where an arbitrary constant velocity traction force drives the harmonic restraint. Here, the velocity of the process is an outcome of the simulation, ruled by the original physical forces and thermal fluctuations at room temperature. In the limit of very small spring constants, the process can be considered fully adiabatic. The choice of the observable to be biased to accelerate the molecular process is crucial to obtain a fast and reliable simulation campaign. In this case, as a CV we chose the distance between the center of mass of the heavy atoms of the N-terminal residue (i.e., Asp237) of the different AP variants facing the mouth of the ribosomal exit channel and a virtual atom in a fixed position 12.5 nm away in the solvent on a line connecting the PTC with the mouth of the exit tunnel. In the ABMD simulations, the system is induced to reduce this distance to zero (Supplementary Fig. 6); the maximal simulation time was set to 100 ns. To set the value of the spring constant, we searched for the highest spring value (fastest simulations) that allowed to see a complete release process without compromising the mechanistic details and differences for the various variants. The force constant (*K*) of the applied harmonic potential was carefully chosen testing a range of *K* between 10^4^ and 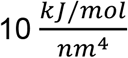 (Supplementary Fig. 9) on the S255A system. The proper choice of *K* allowed us to explore the CV space in a reasonable time and at the same time to adequately sample the intermediate states of the NC release process and discerning the differences between AP behaviors. From the analysis of the test runs, we found that a reasonable choice was 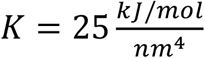. Such a choice was kept constant for all the systems to allow a fair comparison. Performing multiple replicas ABMD simulations allowed us to reduce the variance of the results, as in Gobbo et al.^15^ and Wan et al.^16^, to rank the different variants strength in inducing the stalling of the ribosome. This provided acceptable statistics of 20 replicas for each system, giving a dynamical overview of the atomistic details of the molecular mechanism behind the ribosome arresting induced by human XBPu1, thus complementing the structural and mutational analysis^9,10^.

In the ABMD analysis we defined five metrics (Table 1):

- For the detach time from PTC, we considered the C-terminal residue (CT-Met260) as effectively displaced when the distance covered from its original position was higher than 3.8 Å (the same average distance value between two consecutive Ca);
- Analogously a successfully detached replica had CT-Met260 no more in contact with the PTC;
- the definition of the extraction time is based on the minimal distance between any atom of the AP and any atom of the ribosome; if all these distances are higher than 3.5 Å, then the AP is considered out of the exit channel, and to this time is subtracted the CT-Met260 detach time;
- Analogously a successfully exited replica had no more contacts between the AP and the ribosome;
- The distance covered by the N-terminal residue (Asp237) in CV space is defined as the difference between the initial distance between Asp237 and the target ending point and the final distance between Asp237 and the ending point.

All inter-or intra-molecular contacts were considered lost when their distance was higher than 3.5 Å.

## Acknowledgements

We acknowledge that the research activity herein was carried out using the Italian Institute of Technology (IIT) HPC infrastructure. FDP, SD and AC acknowledge Fondazione Istituto Italiano di Tecnologia for the financial support. Furthermore, FDP acknowledges the CINECA award under the ISCRA initiative (“IsC78_RAP”), for the availability of high performance computing resources. SS kindly acknowledges financial support from the European Research Council under the Horizon 2020 Programme Grant Agreement n. 739964 (“COPMAT”). ML and FP-A were supported by NIH award R35GM122543 and Stanford Data Science Scholar program.

## Author Contribution

FDP designed the research project run simulations, conceived the analysis and analyzed the results, and wrote the paper; SD designed the research project, conceived the simulation protocols and the analysis, analyzed the results and wrote the paper; FP-A,SS and ML wrote the paper; GvH conceived the analysis and analyzed the results, wrote the paper; AC designed the research project and wrote the paper.

