## Supplementary Information for "Probing interplays between human XBP1u translational arrest peptide and 80S ribosome"

### The stalled ribosome: detailed interactions

Here we analyze the resulting interactions along the 1 μs-long (plain) molecular dynamics (MD) simulation of the stalled ribosome with the XBP1u arresting peptide (AP)[1–3] inside the exit tunnel. We start the analysis from the amino acids closest to the P-tRNA up to the terminal part of the ribosome exit tunnel. The same nomenclature of the main text (see Fig. 2) to distinguish AP residues is used: CT- for the C-terminal portion of the AP (residues 260-254); I- for the intermediate part (residues 253-247) and NT- for the N-terminal part (246-237). For the ribosome, we prepend the specific subunit name. The C-terminal CT-Met260 interacts with P-tRNA A76; this interaction was stable for the whole simulation (even without a covalent bond between them that was not included in preparation for the NC release in the Adiabatic Bias MD[4] simulation campaign). In contrast, two non-specific fluctuating contacts were observed with the backbone of 28S-C3909 and with 28S-U4531 (Supplementary Fig. 2). CT-Leu259 interacts with 28S-C4398, 28S-U4452, and 28S-U4531. The first uracil is engaged in an electrostatic interaction with the backbone nitrogen of the leucine, whereas the nucleobases of 28S-C4398 and 28S-U4531 stabilize the leucine side chain (Supplementary Fig. 2). CT-Pro258 is surrounded by 28S-U4531, 28S-G3907, and 28S-A3908 (Supplementary Fig. 2). The latter additionally stabilizes the CT-Lys257 backbone, side chain of which stacks with 28S-U4532 via a pi-cation interaction. This stacking interaction, combined with the pi-cation between the positively charged ε-amino group and the sugar of 28S-U4531, contributes to limit the movement of these PTC bases (Supplementary Fig. 2). Moreover, these C-terminal residues, in particular CT-Leu259, partially fill the space that the incoming loaded A-tRNA should occupy; indeed, the resulting complex network of interactions rigidify the structure around the PTC, preventing the ribosome cycle to move on.

CT-Ala255 interactions with 28S-U4532 and 28S-U4555 were initially absent but early established and stable, even if non-specific (Supplementary Fig. 3). CT-Pro254 was in close contact with 28S-U3644, 28S-A3908, 28S-U4552, and 28S-U4555, forming a partial stacking (Supplementary Fig. 3). 28S-U4552 is also involved in a very stable hydrogen bond with I-Gln253 reporting few fluctuations in the simulated time-scale (Supplementary Fig. 3). I-His252 showed a transient and unusual T-shaped pi-stacking interaction with uL22-His133 ribosomal protein (Supplementary Fig. 3).

The charged side-chain position of I-Arg251 was initially stabilized by a salt bridge with the phosphate of 28S-A4388; after 50 ns the salt bridge switched to the phosphate of 28S-A3908, gradually gaining a new stable configuration (Supplementary Fig. 3). This nucleobase stabilizes I-Gly250 via an H-bond that becomes transient and discontinuous in the second half of the simulation (Supplementary Fig. 3). I-Trp249 (also mentioned in the main text) side chain was persistently flanked by 28S-U4557 and at the same time obtained a transient H-bond with uL4-Arg71; this last residue lost its interaction with the backbone of NT-Pro243 (Supplementary Fig. 4).

Even if in the starting structure[3] I-Gln248, I-Cys247, and NT-Leu246 were not bearing any particular interaction (except a very unstable contact between I-Cys247 and 28S-U4555), they found pretty early some interacting partners namely uL22-Arg135, uL22-Gly134, uL22-His133, respectively (Supplementary Fig. 4). We found a very stable π-cation between NT-Phe245 and uL4-Arg71. In contrast, the initial contacts of this arginine with NT-Pro244 and NT-Pro243 (due to a rotation of 180° of the arginine $\chi_1$ dihedral angle) were almost completely lost. The π–π stacking between NT-Tyr241 and 28S-C2794 was preserved almost till the end of the simulation, becoming unstable after ~870 ns (Supplementary Fig. 4); this gave another indication about the weak nature of the contacts involving the AP amino acids that are close to the mouth of the exit channel. At variance of NT-Val239 that had no persistent contacts to mention, both NT-Pro240 and NT-Pro238 interacted with uL22-Arg128 side chain (Supplementary Fig. 4). Moreover, NT-Pro238 was also locked between the backbone of 28S-C368 and 28S-C2794 rRNA (Supplementary Fig. 4). It is finally worth mentioning that the N-terminal end of NT-Asp237 established an H-bond with the phosphate of 28S-C368 (Supplementary Fig. 4).

**Extraction of AP variants from the exit tunnel: detailed interactions**

In the C247K/S255A ABMD simulations (Supplementary Fig. 6e/f), the initial interactions lasted all the way long (Supplementary Fig. 8e). This further confirmed that C247K/S255A variant was stacked inside the channel, mainly due to the positively charged lysine that was introduced with the mutation at residue 247. This is somewhat in accordance with the conclusions in Nissley et al.[5], in which nascent chains carrying several positive charges have been found to be ejected out of the *E. coli* ribosome exit tunnel slowly.

In the simulations with the C247S/P254C/S255A variant (Supplementary Fig. 6d), a clear pattern common to all replicas appeared: the complete solvation of the AP was prevented by prolonged interactions (more than 35 ns) that retained the C-terminal residues inside the exit channel (Supplementary Fig. 8d). These interactions were CT-Cys254 with 28S-A1600 and 28S-C2794, CT-Ala255 with uL22-Arg135, CT-Trp256 with uL4-Arg71.

Conversely, in both the S255A and the W256A variants (Supplementary Fig. 6b and c, respectively), CT-Cys254 and CT-Ala255/Ser255 could not find any stable interaction along the tunnel, thus not preventing their release into the solvent (Supplementary Fig 8b and c). Even if the transient inter-molecular interactions reported for the mutated AP residues of the variants were different, in S255A (Supplementary Fig 8b), all the key contacts with the ribosome were lost after ~30 ns of Adiabatic Bias MD simulation[4]. In contrast, in W256A (Supplementary Fig 8c), this happened at >40 ns. Eventually, such a difference resulted in an increased total release time for the W256A variant relative to S255A (Table 1).

The behavior of the WT ABMD replicas was globally very similar to the S255A variant one. The nascent chain was released faster than any of the variants from the ribosome exit channel into the solvent (Table 1), even if its detachment was slightly slower than the W256A variant one. The pattern of interactions established by the WT residue interested by mutation in the different variants (i.e. 247, 254, 255, 256) was a kind of mix between the S255A and W256A (Supplementary Fig. 8a): NT-Cys247 and NT-W256 carried almost the same interactions as in S255A; CT-Pro254 interacted with 28S-A3908, 28S-U4555, uL22-Gly134 and uL4-His85 like in S255A, and with 28S-G4527, 28S-U4556 and 28S-C368 like in W256A; CT-Ser255 pattern of interactions was identical to the one in W256A, but the WT interactions lasted on average quite less than the ones in the W256A replicas.

We also calculated the average distance (D) covered by the WT and the different AP variants (Table 1). We obtained D = 12.5 nm ± 0.00 in both the WT and the S255A variant as they always successfully reached the end point in all the 20 replicas as defined in the ABMD protocol. For W256A, not all the replicas (15/20) reached the end point, thus $D_{W256A}$ = 11.7 nm ± 0.8, slightly below the previously mentioned distances. For the triple mutant, C247S/P254C/S255A, we calculated $D_{C247S/P254C/S255A}$ = 7.4 nm ± 2.6 with not even one replica in which the nascent chain resulted fully ejected in the solvent even if

17 of them detached from the PTC. For C247K/S255A, the distance covered was only $D_{C247K/S255A}$ = 0.47 nm ± 0.01, thus no detaching from PTC nor nascent chain ejection was possible.

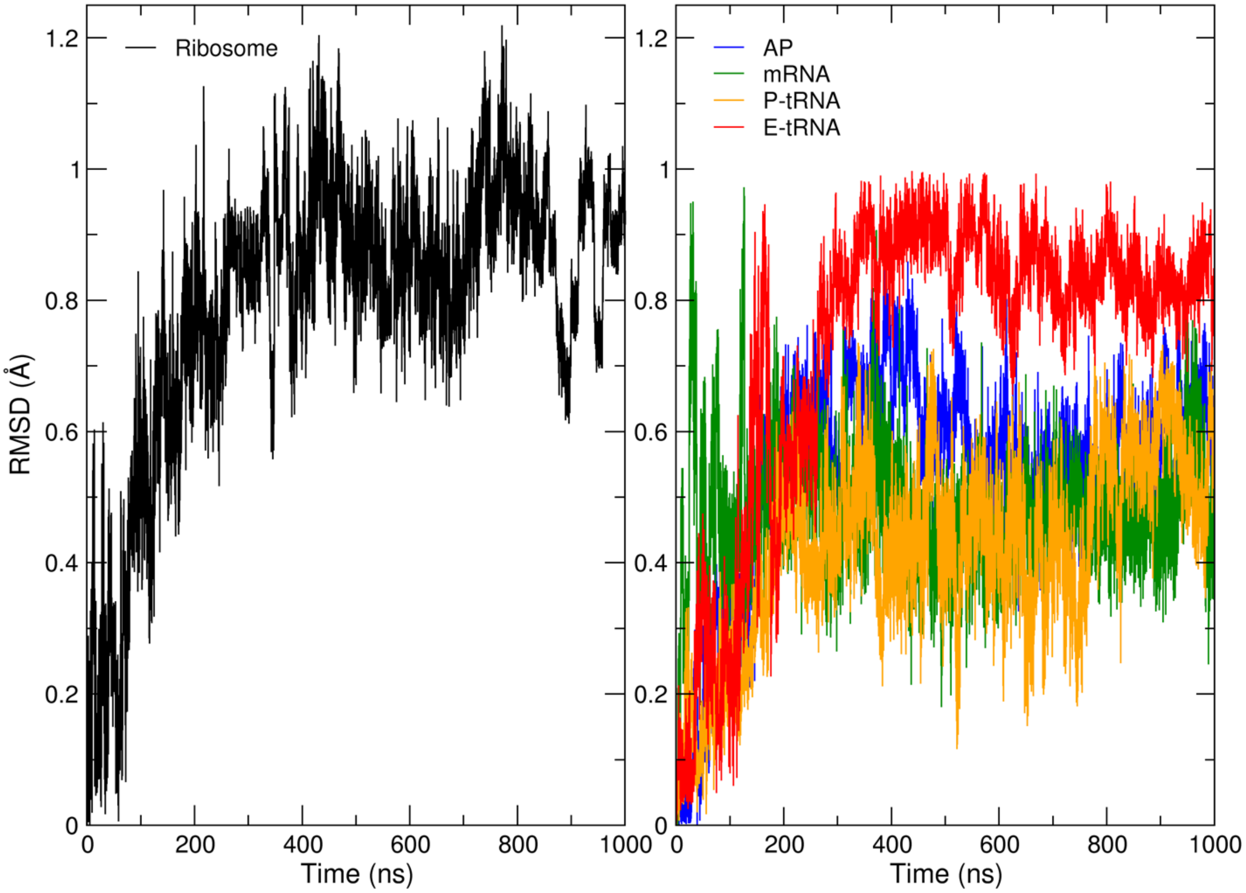


**Supplementary Fig. 1.** Root Mean Square Deviation of the 1μs equilibrium simulation. a) RMSD of the sugar-phosphate and protein backbone atoms of the whole Ribosome. b) RMSD of the backbone of AP (blue). mRNA (green). P-tRNA (Orange). E-tRNA (red).

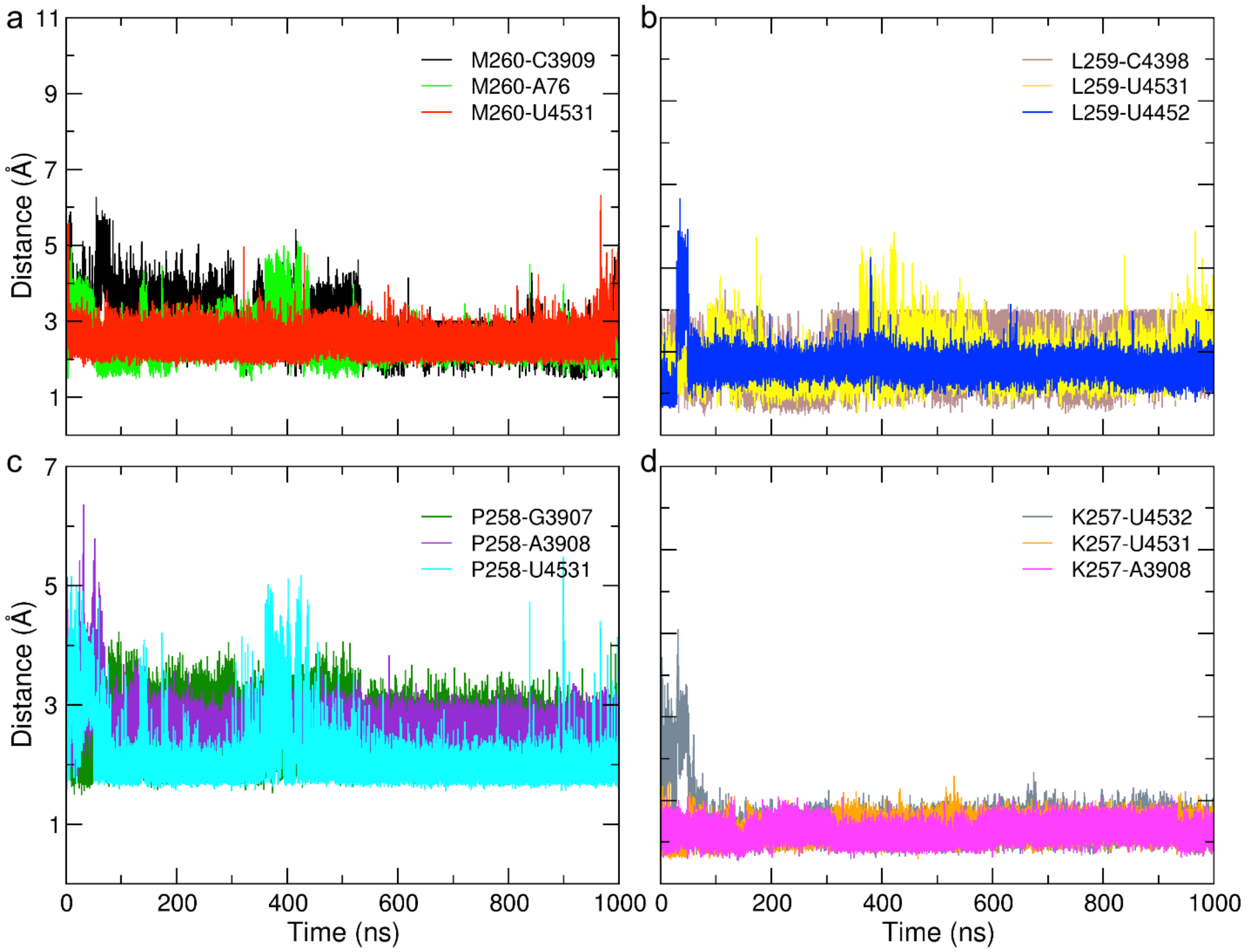


**Supplementary Fig. 2.** Intermolecular contacts between AP and ribosome over 1 μs equilibrium simulation. In each graph, the following AP residues interacting with their partners can be found a) M260; b) L259; c) P258; d) K257. A76 in panel “a” is the 3’ nucleobase of the P-site tRNA.

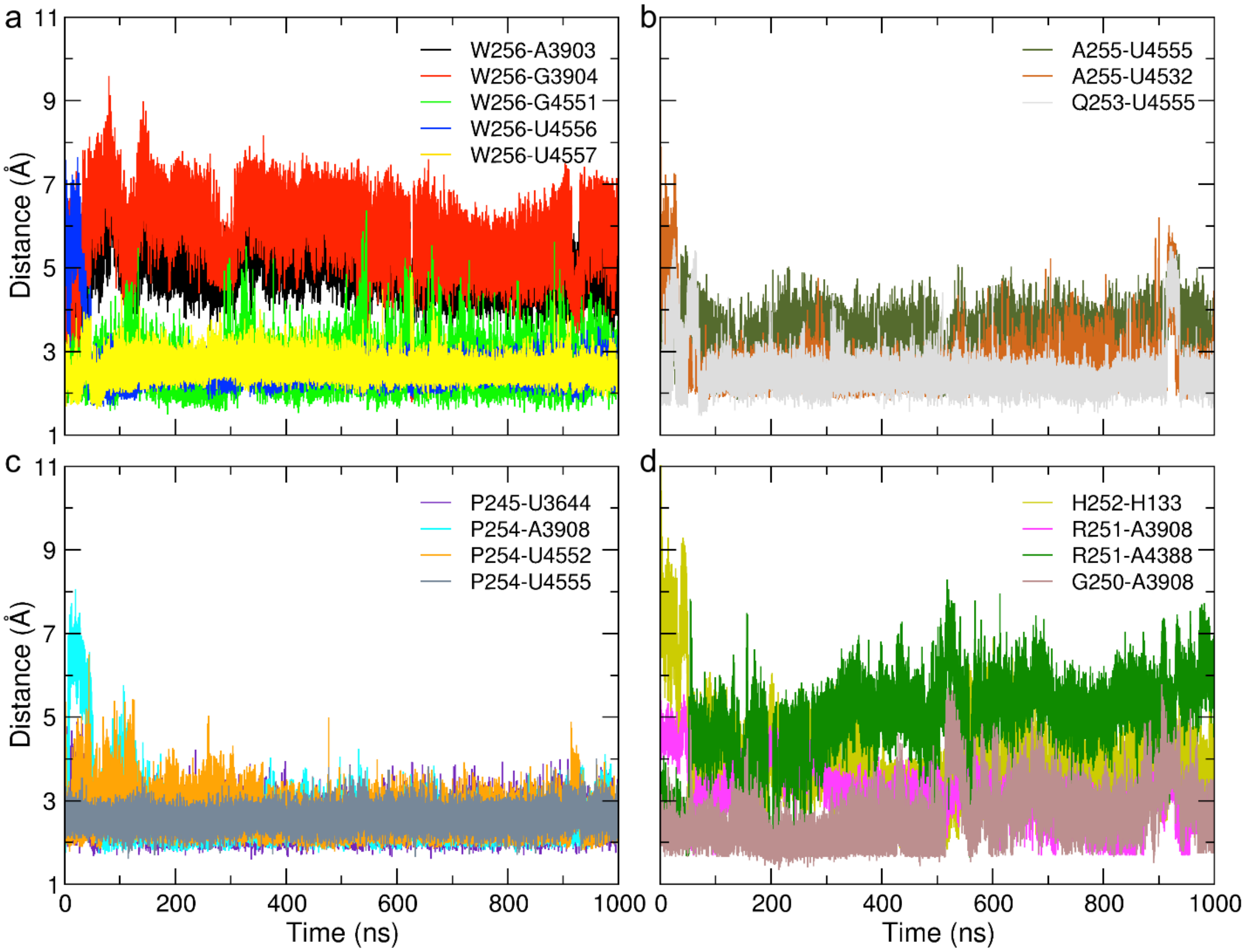


**Supplementary Fig. 3.** Intermolecular contacts between AP and ribosome over 1 μs equilibrium simulation. In each graph, the following AP residues, interacting with their partners, can be found a) W256; b) A255 and Q253; c) P254; d) H252. R251 and G250.

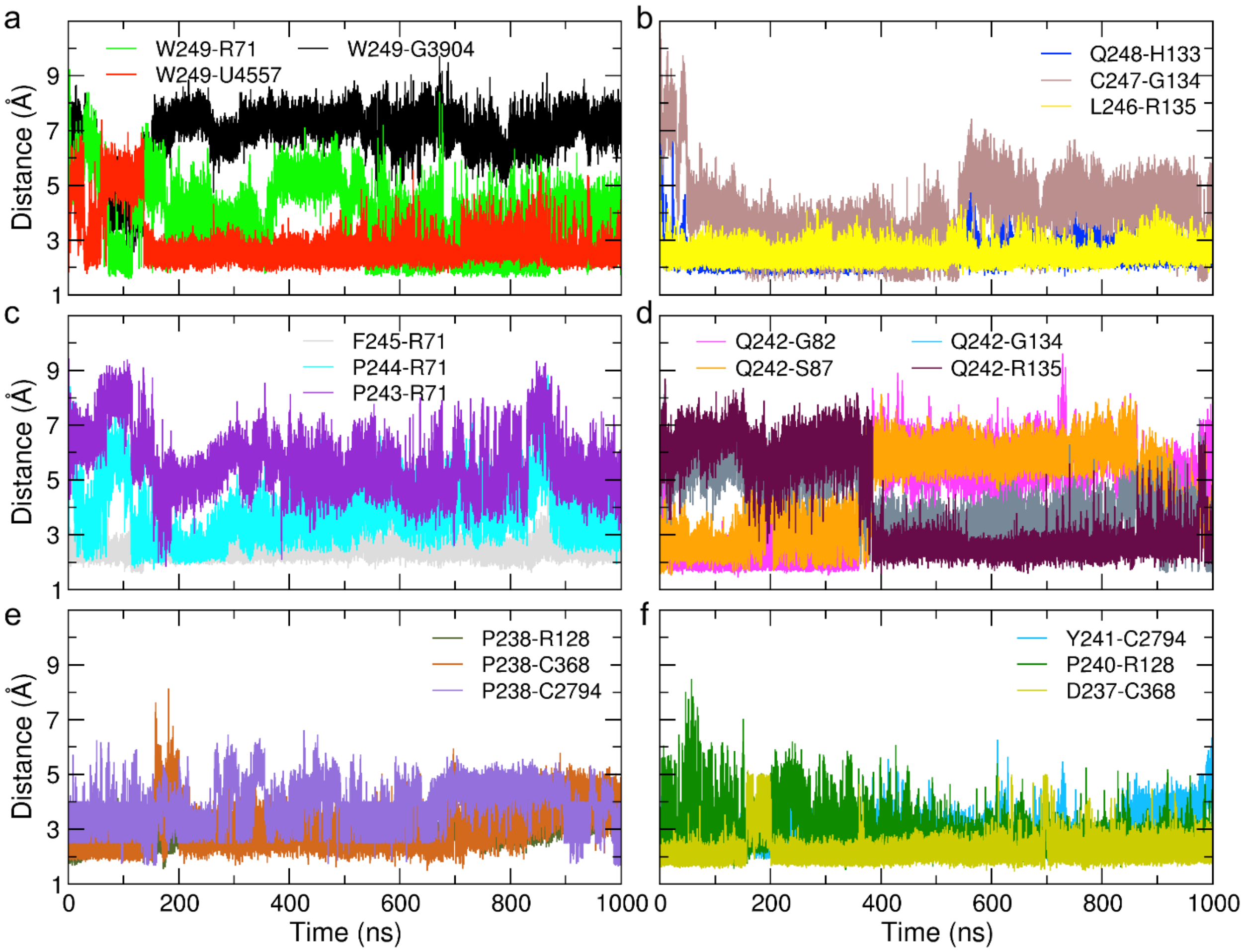


**Supplementary Fig. 4.** Intermolecular contacts between AP and ribosome over 1 μs equilibrium simulation. In each graph, the following AP residues, interacting with their partners, can be found a) W249; b) Q248, C247, and L246; c) F245, P244, P243; d) Q242; e) P238; f) Y241, P240, and D237.

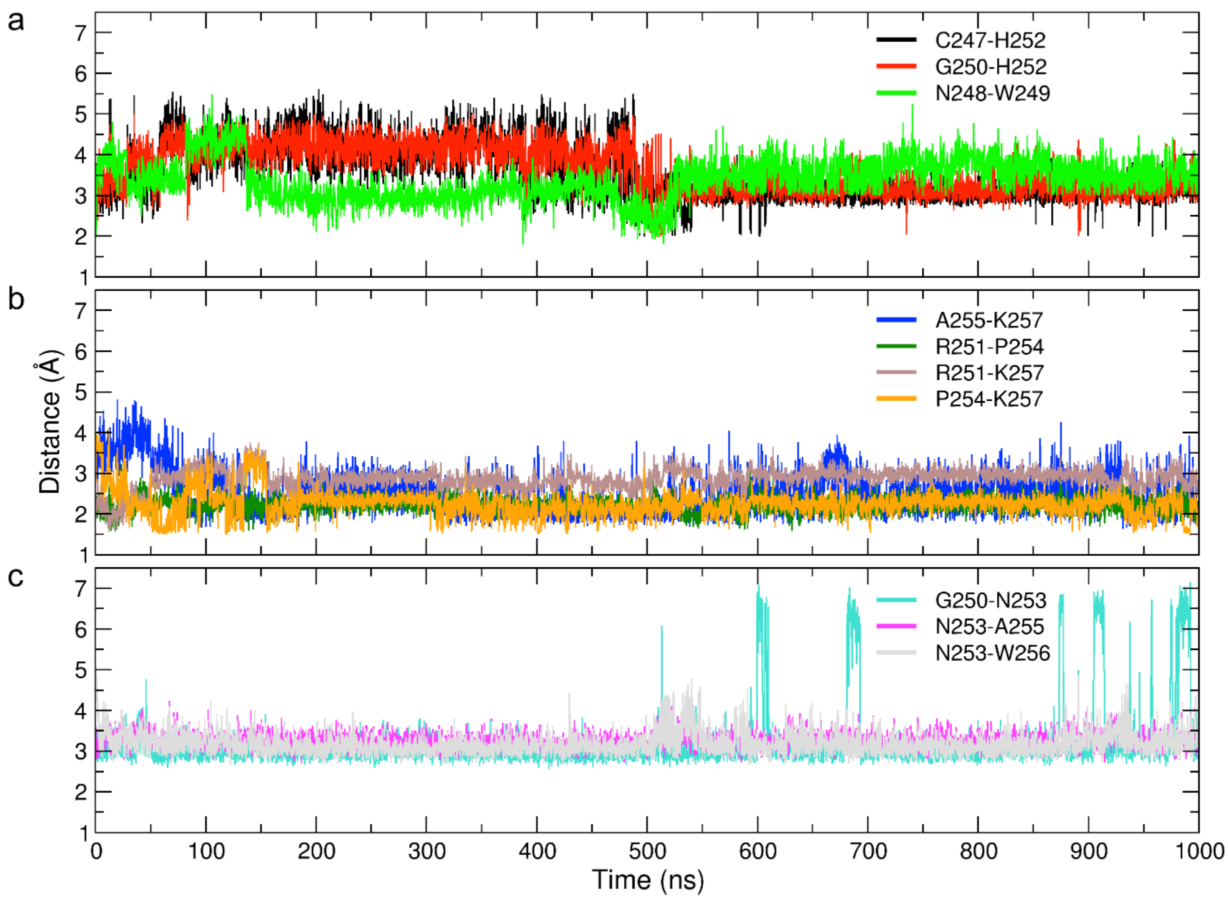


**Supplementary Fig. 5.** AP intramolecular contacts over 1 μs of MD. The three networks, as described in the text are split in the 3 panels: a) C247, Q248 and W249 (CN1); b) G250, Q253, and W256 (CN2); c) R251, P254, A255, and K257 (CN3).

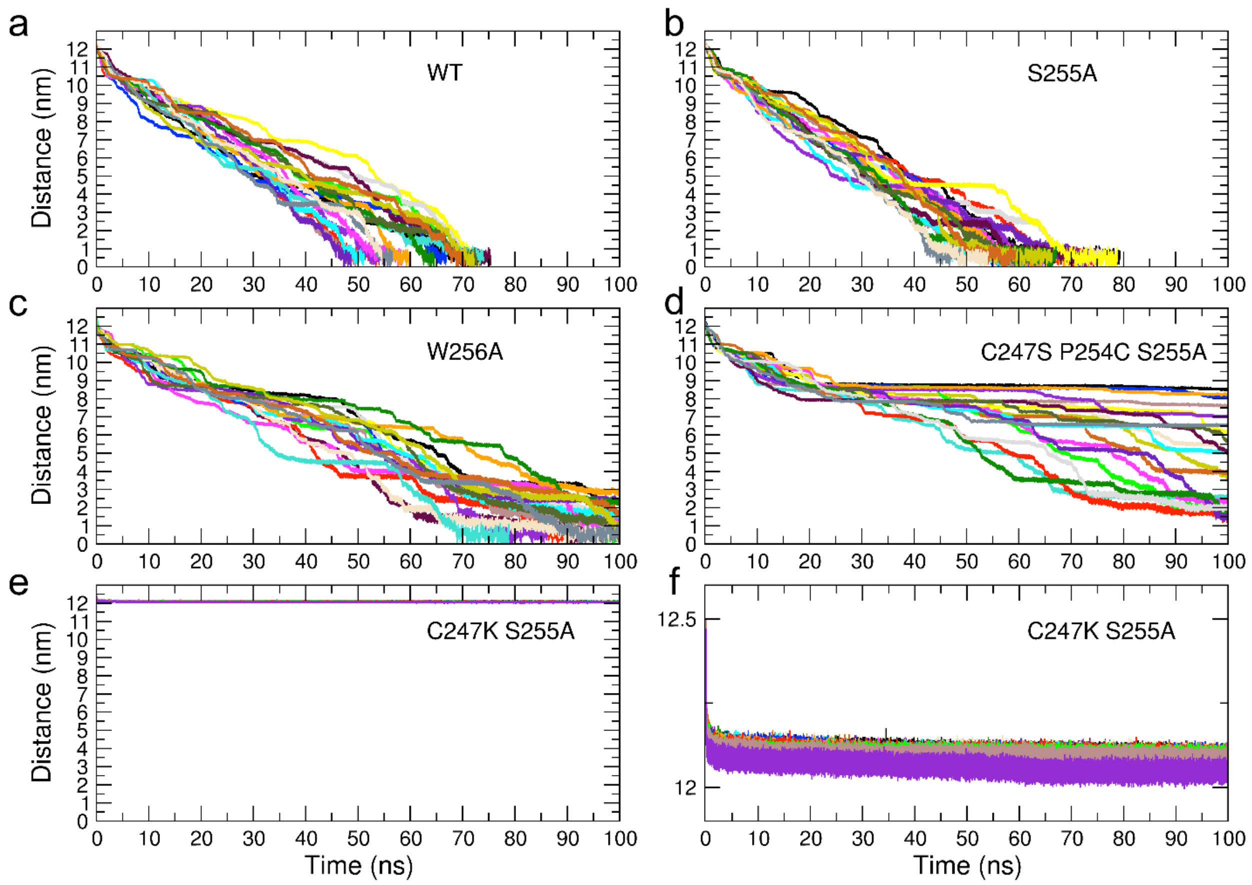


**Supplementary Fig. 6.** Resulting CV time-course from the 20 independent ABMD replicas for each AP variant. In the e) panel, C247K/S255A variant, using the same y-axis interval of the other plots for comparison purpose, the 20 curves appear completely overlapped, thus, its zoom is shown in the f) panel for clarity.

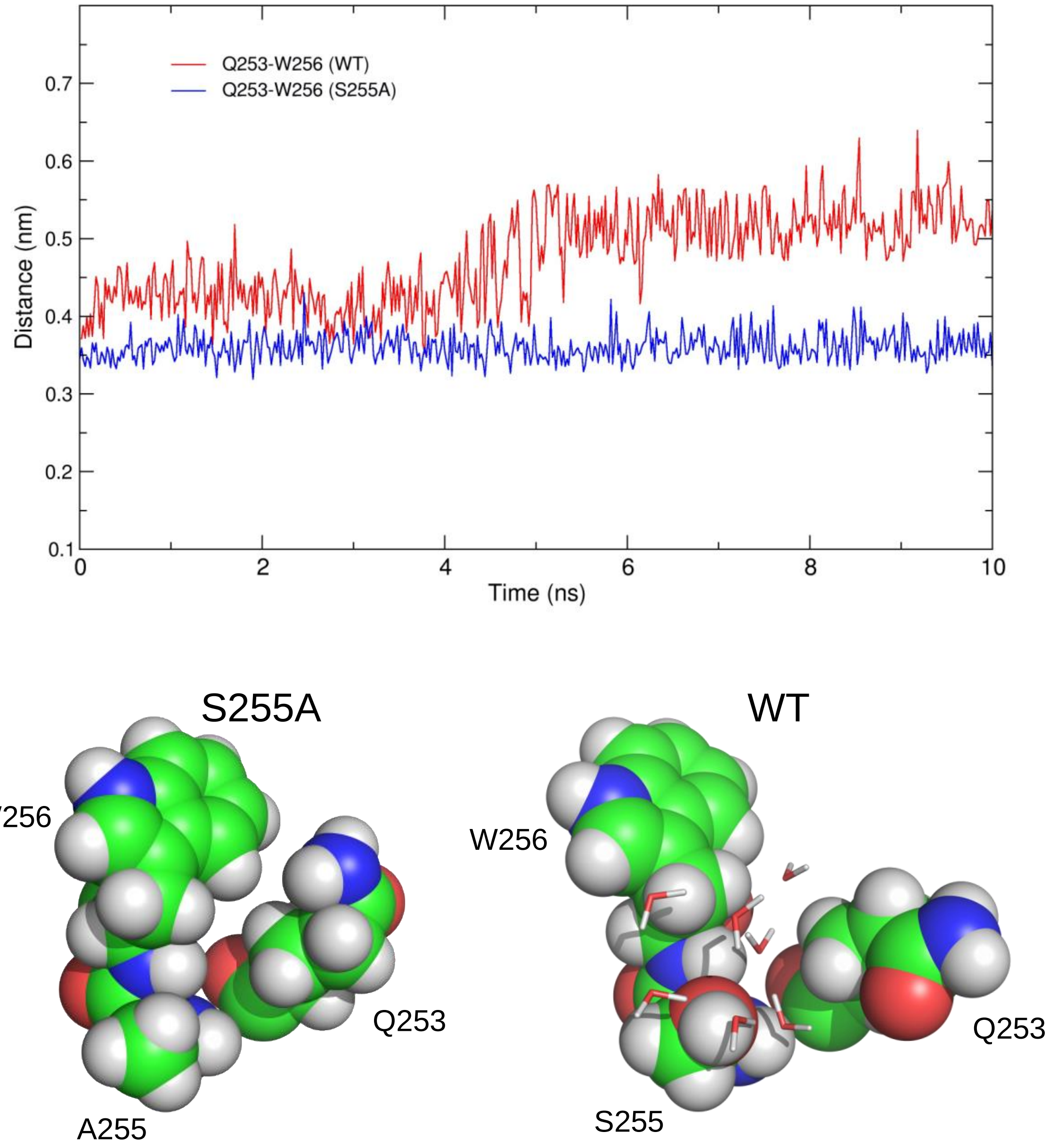


**Supplementary Figure 7.** Comparative Q253-W256 interaction time-course from the WT and S255A variant 10 ns equilibration simulations. The molecular models show the close-up on the Q253-W256 interaction in the presence of CT-A255 (left, S255A variant) and of CT-S255 (right, WT). In the latter case, the presence of several water molecules solvating serine side chain prevents an important packing interaction between Gln253 and Trp256 seen in S255A variant.

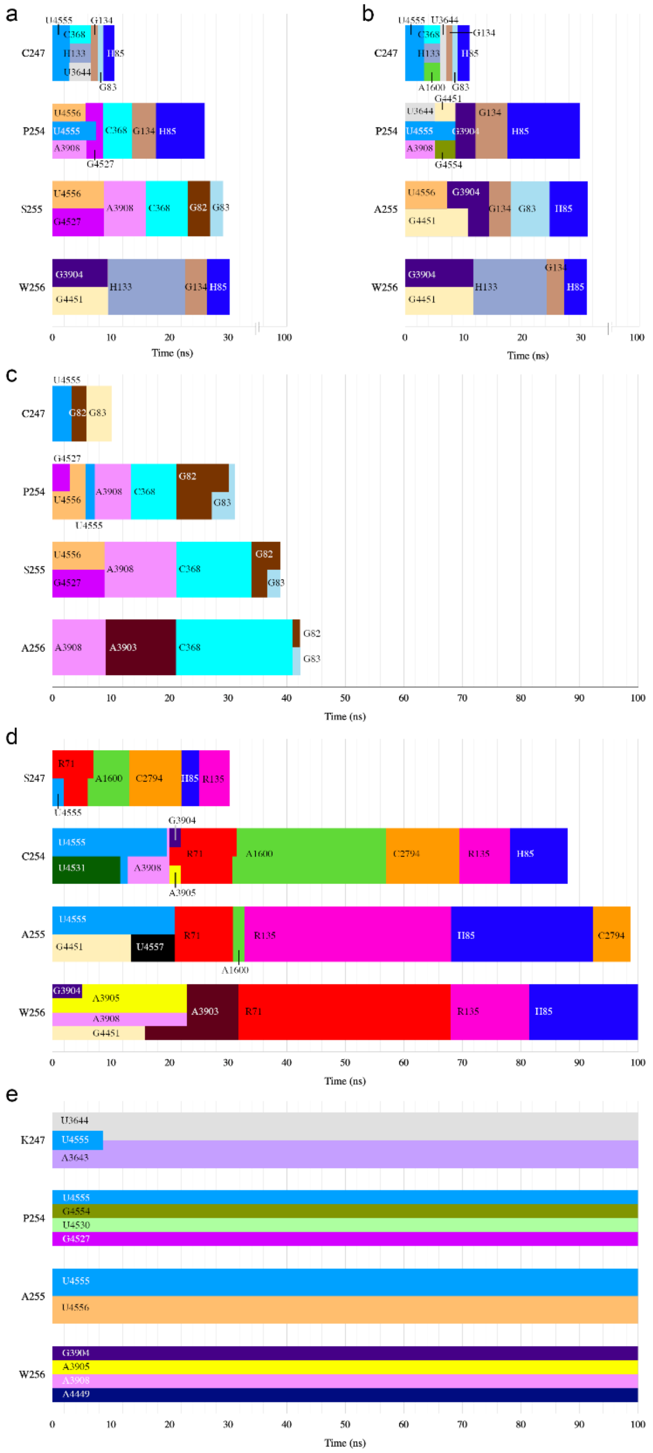


**Supplementary Fig. 8.** Conserved interactions in the 100 ns ABMD simulations of the mutated residues (numbers 247, 254, 255 and 256) in the four variants. The early part of each graph, representing the trajectories up to the point when individual residues detach from the PTC, are the same as in Fig. 5 in the main text, while the remaining part represents the extraction phase after the detachment of Met260 from the PTC, in which the AP is pulled out of the ribosome. a) WT, b) S255A, c) W256A, d) C247S/P254C/S255A, e) C247K/S255A. For each residue the colored bar represents the range of time (in nanoseconds) in which the AP was interacting with the specific residue/nucleobase of the ribosome. The end of the bar represents the first time instant for which in at least one of the simulations the interaction was lost (the bar width does not encode any data; it is just for convenience of graphical representation).

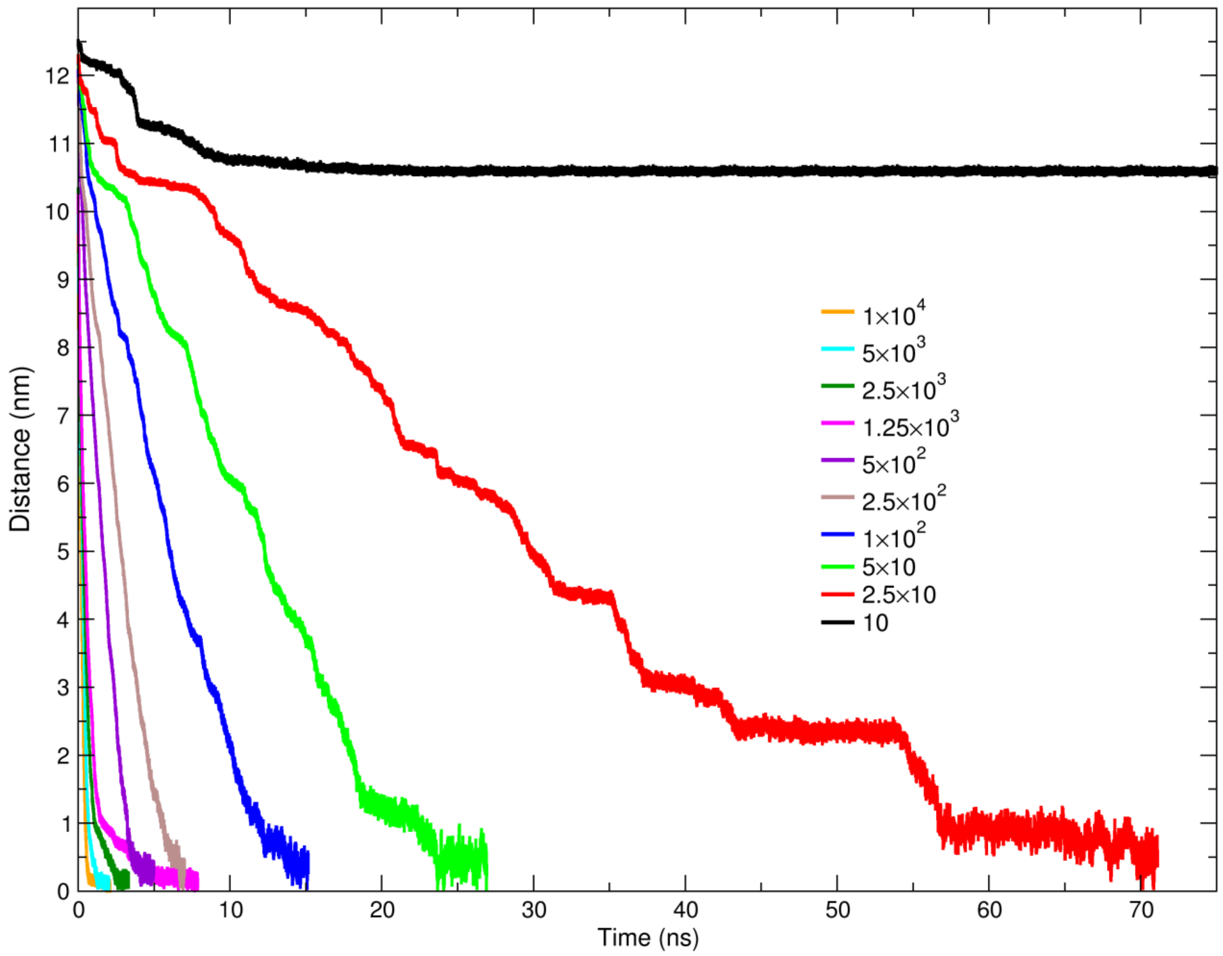


**Supplementary Fig. 9.** Resulting CV time-course from nine trials of different force constant ($K$) of the ABMD harmonic potential spanning from 10 to $10^4$ $\frac{kJ/mol}{nm^4}$. In the second simulative campaign, to properly sample the intermediate states of the APs exit process, $K$ = 25 $\frac{kJ/mol}{nm^4}$ was set up.

**Supplementary Table 1.** Intermolecular AP-ribosome contacts along the equilibrium 1 μs long all-atom MD simulation.

| AP residue | Ribosomal bases/residue | Persistency | Behavior |
|---|---|---|---|
| M260 | A76 (P-tRNA) | .76 | Half fluctuating/half stable |
| " | C3909 (28S rRNA) | .67 | Bimodal |
| " | U4531 (28S rRNA) | .99 | Stable |
| L259 | U4452 (28S rRNA) | .93 | Stable |
| " | C4398 (28S rRNA) | .69 | Fluctuating |
| " | U4531 | .81 | Some fluctuations |
| P258 | G3907 (28S rRNA) | .90 | Few fluctuations |
| " | A3908 (28S rRNA) | .92 | Few initial fluctuation |
| " | U4531 | .92 | Few fluctuations |
| K257 | A3908 | 1.0 | Stable |
| " | U4531 | 1.0 | Stable |
| " | U4532 (28S rRNA) | .94 | Stable after 50 ns |
| W256 | A3903 (28S rRNA) | .03 | Lost at 35 ns |
| " | G3904 (28S rRNA) | .01 | Lost in few ns |
| " | G4551 (28S rRNA) | .96 | Stable |
| " | U4556 (28S rRNA) | .96 | Gained in few ns |
| " | U4557 (28S rRNA) | .99 | Stable |
| A255 | U4532 | .90 | Gained at 35 ns |
| " | U4555 (28S rRNA) | .30 | Fluctuating a lot |

| | | | |
|---|---|---|---|
| “ | U4552 (28S rRNA) | .97 | Stable |
| P254 | U3644 (28S rRNA) | .93 | Stable |
| “ | A3908 | .90 | Stable |
| “ | U4555 | .85 | Initial fluctuation |
| Q253 | U4555 | .93 | Few fluctuations |
| H252 | H133 (uL22) | .32 | Gained at 50ns/transient |
| “ | A4388 (28S rRNA) | .08 | Lost after 50 ns |
| R251 | A3908 | .71 | Gained at 60ns/fluctuating |
| G250 | A3908 | .76 | Transient after 500ns |
| W249 | G3904 | .01 | Lost in few ns |
| “ | U4557 | .87 | Quite Stable |
| “ | R71 (uL4) | .42 | Gained at 70 ns/transient |
| Q248 | H133 (uL22) | .94 | Stably gained at 25ns |
| C247 | U4555 | .08 | Definitely lost after 100ns |
| C247 | G134 (uL22) | .48 | Gained at 50ns/transient |
| L246 | R135 (uL22) | .94 | Stably Gained in few ns |
| F245 | R71 | .97 | Stable |
| P244 | R71 | .32 | Highly Fluctuating |
| P243 | R71 | .02 | Lost in few ns |
| Q242 | G82 (uL4) | .34 | Lost at 400ns |
| “ | S87 (uL4) | .31 | Lost at 400ns |
| “ | G134 | .40 | Gained at 400ns |
| “ | R135 | .59 | Gained at 400ns |

| | | | |
|---|---|---|---|
| Y241 | C2794 (28S rRNA) | .88 | Stable till 870 ns |
| P240 | R128 (uL22) | .87 | Almost stable |
| P238 | R128 | .59 | Bimodal |
| “ | C368 (28S rRNA) | .85 | Stable till 800ns |
| “ | C2794 | .56 | Bimodal |
| D237 | C368 | .94 | Barely Fluctuating |

**Supplementary Table 2.** Intramolecular AP-AP contacts along the equilibrium 1 μs long all-atom MD simulation.

| AP residue 1 | AP residue 2 | Persistency | Behavior |
|---|---|---|---|
| C247 | H252 | .56 | First half fluctuating/<br>second half very stable |
| Q248 | W249 | .89 | Short fluctuations after 100 ns |
| G250 | H252 | .55 | First half fluctuating/<br>second half very stable |
| R251 | P254 | 1.0 | Stable |
| R251 | K257 | .99 | Stable |
| P254 | K257 | .88 | Few initial fluctuations |
| A255 | K257 | .80 | Initially fluctuating,<br>quite stable after 150 ns |
| Q253 | G250 | .92 | Short fluctuations after 500 ns |
| Q253 | A255 | .98 | Stable |
| Q253 | W256 | .97 | Stable |

**Supplementary Table 3.** Original inter- and intra-molecular contacts average time-loss (in ns), and associated standard error, along the out-of-equilibrium 100 ns ABMD[4] simulations (as plotted in Fig. 4a). An asterisk means that the corresponding residue never lost its contacts.

| AP residue | WT | S255A | W256A | C247S/P254C/S255A | C247K/S255A |
|---|---|---|---|---|---|
| 237 | 0.20 ± 0.01 | 0.22 ± 0.01 | 0.23 ± 0.01 | 0.28 ± 0.01 | 0.26 ± 0.01 |
| 238 | 0.21 ± 0.01 | 0.25 ± 0.01 | 0.25 ± 0.01 | 0.30 ± 0.01 | 0.30 ± 0.01 |
| 239 | 0.22 ± 0.02 | 0.25 ± 0.01 | 0.26 ± 0.01 | 0.31 ± 0.01 | 0.35 ± 0.01 |
| 240 | 0.95 ± 0.04 | 1.03 ± 0.03 | 1.11 ± 0.04 | 1.25 ± 0.05 | 6.12 ± 4.72 |
| 241 | 1.33 ± 0.02 | 1.59 ± 0.04 | 1.61 ± 0.05 | 1.72 ± 0.07 | 25.18 ± 10.07 |
| 242 | 1.38 ± 0.04 | 1.60 ± 0.05 | 1.64 ± 0.05 | 1.75 ± 0.10 | * |
| 243 | 1.53 ± 0.02 | 1.78 ± 0.04 | 1.77 ± 0.05 | 1.83 ± 0.11 | * |
| 244 | 1.55 ± 0.05 | 1.80 ± 0.06 | 1.79 ± 0.06 | 1.99 ± 0.19 | * |
| 245 | 1.60 ± 0.06 | 1.85 ± 0.05 | 1.83 ± 0.10 | 2.02 ± 0.22 | * |
| 246 | 1.79 ± 0.11 | 2.05 ± 0.13 | 2.01 ± 0.21 | 3.46 ± 0.42 | * |
| 247 | 2.95 ± 0.25 | 3.22 ± 0.25 | 3.19 ± 0.30 | 7.13 ± 0.38 | * |
| 248 | 2.99 ± 0.36 | 3.25 ± 0.37 | 3.25 ± 0.34 | 8.11 ± 0.43 | * |
| 249 | 4.30 ± 0.42 | 4.57 ± 0.40 | 4.55 ± 0.51 | 9.38 ± 0.59 | * |
| 250 | 4.66 ± 0.39 | 4.96 ± 0.36 | 4.90 ± 0.47 | 11.05 ± 1.23 | * |
| 251 | 5.29 ± 0.60 | 5.67 ± 0.59 | 5.58 ± 0.70 | 12.29 ± 1.18 | * |
| 252 | 5.33 ± 0.65 | 5.67 ± 0.61 | 5.60 ± 0.66 | 13.97 ± 1.46 | * |
| 253 | 5.87 ± 0.69 | 6.45 ± 0.69 | 6.17 ± 0.72 | 15.06 ± 2.09 | * |
| 254 | 7.72 ± 0.80 | 8.66 ± 0.85 | 8.10 ± 0.81 | 19.59 ± 3.13 | * |

| | | | | | |
|---|---|---|---|---|---|
| 255 | 8.81 ± 0.88 | 10.74 ± 0.92 | 8.89 ± 0.98 | 20.84 ± 2.97 | * |
| 256 | 9.55 ± 0.91 | 11.59 ± 0.88 | 9.15 ± 0.92 | 23.03 ± 3.56 | * |
| 257 | 10.53 ± 0.78 | 12.48 ± 0.81 | 10.47 ± 1.03 | 24.15 ± 5.02 | * |
| 258 | 11.49 ± 0.90 | 13.31 ± 0.97 | 11.34 ± 1.10 | 25.31 ± 4.86 | * |
| 259 | 12.55 ± 0.92 | 13.67 ± 1.01 | 12.45 ± 1.14 | 26.14 ± 5.23 | * |
| 260 | 13.16 ± 0.89 | 13.99 ± 0.94 | 13.02 ± 1.09 | 27.12 ± 5.61 | * |

**Supplementary Table 4.** Average time-intervals <$\Delta t^i$> (in ns), and associated standard error, required to detach each AP residue after the previous one has detached during the 100 ns ABMD[4] simulations (as plotted in Fig. 4b). A star symbol means that the corresponding residue never detached.

| AP residue | WT | S255A | W256A | C247S/P254C/S255A | C247K/S255A |
|---|---|---|---|---|---|
| 237 | 0.20 ± 0.01 | 0.22 ± 0.01 | 0.23 ± 0.01 | 0.28 ± 0.01 | 0.26 ± 0.01 |
| 238 | 0.01 ± 0.01 | 0.03 ± 0.01 | 0.02 ± 0.01 | 0.02 ± 0.01 | 0.30 ± 0.01 |
| 239 | 0.01 ± 0.01 | 0.01 ± 0.01 | 0.01 ± 0.01 | 0.01 ± 0.01 | 0.35 ± 0.01 |
| 240 | 0.73 ± 0.04 | 0.77 ± 0.02 | 0.85 ± 0.03 | 0.94 ± 0.05 | 5.77 ± 3.14 |
| 241 | 0.58 ± 0.02 | 0.56 ± 0.05 | 0.51 ± 0.03 | 0.47 ± 0.07 | 19.06 ± 7.12 |
| 242 | 0.05 ± 0.01 | 0.01 ± 0.01 | 0.03 ± 0.03 | 0.03 ± 0.02 | * |
| 243 | 0.16 ± 0.02 | 0.18 ± 0.03 | 0.13 ± 0.03 | 0.08 ± 0.03 | * |
| 244 | 0.02 ± 0.01 | 0.02 ± 0.01 | 0.02 ± 0.01 | 0.16 ± 0.04 | * |
| 245 | 0.05 ± 0.02 | 0.05 ± 0.04 | 0.04 ± 0.01 | 0.12 ± 0.02 | * |
| 246 | 0.20 ± 0.05 | 0.20 ± 0.09 | 0.18 ± 0.04 | 1.44 ± 0.12 | * |
| 247 | 1.17 ± 0.08 | 1.17 ± 0.07 | 1.18 ± 0.11 | 3.67 ± 0.27 | * |
| 248 | 0.04 ± 0.01 | 0.03 ± 0.02 | 0.06 ± 0.04 | 0.98 ± 0.30 | * |
| 249 | 1.31 ± 0.22 | 1.32 ± 0.18 | 1.30 ± 0.36 | 1.27 ± 0.42 | * |
| 250 | 0.37 ± 0.04 | 0.39 ± 0.05 | 0.35 ± 0.03 | 1.67 ± 0.46 | * |
| 251 | 0.64 ± 0.12 | 0.71 ± 0.11 | 0.68 ± 0.15 | 1.24 ± 0.28 | * |
| 252 | 0.03 ± 0.01 | 0.01 ± 0.01 | 0.02 ± 0.01 | 1.68 ± 0.36 | * |
| 253 | 0.54 ± 0.13 | 0.77 ± 0.08 | 0.57 ± 0.10 | 1.09 ± 0.27 | * |
| 254 | 1.86 ± 0.16 | 2.21 ± 0.20 | 1.93 ± 0.17 | 4.53 ± 0.81 | * |

| | | | | | |
|---|---|---|---|---|---|
| 255 | 1.11 ± 0.16 | 2.08 ± 0.18 | 0.79 ± 0.19 | 1.25 ± 0.19 | * |
| 256 | 0.84 ± 0.12 | 0.85 ± 0.12 | 0.26 ± 0.10 | 2.19 ± 0.52 | * |
| 257 | 0.93 ± 0.18 | 0.89 ± 0.17 | 1.30 ± 0.22 | 1.12 ± 0.22 | * |
| 258 | 0.92 ± 0.22 | 0.83 ± 0.18 | 0.85 ± 0.20 | 1.16 ± 0.26 | * |
| 259 | 0.86 ± 0.12 | 0.36 ± 0.06 | 1.11 ± 0.14 | 0.83 ± 0.13 | * |
| 260 | 0.59 ± 0.10 | 0.32 ± 0.05 | 0.58 ± 0.09 | 0.98 ± 0.11 | * |
